# Obstacles Regulate Membrane Tension Propagation to Enable Localized Mechanotransduction

**DOI:** 10.1101/2025.01.14.632796

**Authors:** Frederic Català-Castro, Mayte Bonilla-Quintana, Neus Sanfeliu-Cerdán, Padmini Rangamani, Michael Krieg

**Affiliations:** ICFO - Institut de Ciències Fotòniques, Castelldefels, The Barcelona Institute of Science and Technology, Barcelona, Spain; Department of Mechanical and Aerospace Engineering, University of California San Diego, La Jolla, USA; Department of Pharmacology, School of Medicine, University of California San Diego, La Jolla, USA

## Abstract

Forces applied to cellular membranes lead to transient membrane tension gradients. The way membrane tension propagates away from the stimulus site into the membrane reservoir is a key property in cellular adaptation. However, it remains unclear how tension propagation in membranes is regulated and how it depends on the cell type. Here, we investigate plasma membrane tension propagation in cultured *Caenorhabditis elegans* mechanosensory neurons. We show that tension propagation travels quickly and is restricted to a particular distance in the neurites — projections from the cell body of a neuron. A biophysical model of tension propagation suggests that periodic obstacle density and arrangement play key roles in controlling the propagation of mechanical information. Our experiments show that tension propagation is strongly dependent on the intact actin and microtubule cytoskeleton, whereas membrane lipid properties have minimal impact. In particular, the organization of the *α/β*-spectrin network and the MEC-2 stomatin condensates in periodic scaffold act as barriers to tension propagation, limiting the spread of tension. Our findings suggest that restricting membrane tension propagation in space and time enables precise localized signaling, allows a single neuron to process mechanical signals in multiple distinct domains, thus expanding its computational capacity.

## 1 Main text

The transmission of mechanical information through the cytoskeleton, organelles, and plasma membrane is vital for the organization of cells and mechanotransduction[1, 2]. Plasma membrane mechanics plays an important role in the cellular response to external forces and determines cell shape [3], signaling [4], and metabolism [5]. Mechanical properties of the membrane are characterized by an in-plane membrane elasticity, an elastic restoring force that resists bending deformations, a viscosity that resists flow and, originating from its coupling to the cell cortex, an effective membrane tension [6]. Furthermore, it has been extensively reported that local tension gradients, due to a large membrane reservoir or the fluid character of the lipid bilayer, rapidly relax in metazoan cells [7, 8, 9, 10, 11, 12] but may be heterogeneous along the contour of the cell [13, 14, 15, 16]. Thus, the presence of such a lipid reservoir implies that the membrane cannot support or store mechanical stresses on timescales longer than its viscous relaxation time, resulting in a constant membrane tension [7]. Due to this, it is believed that a local increase in tension generated by an application of local forces (due to an external tether extrusion [7] or an internal actin protrusion [17]) is rapidly propagated across the entire membrane surface. This idea has placed membrane tension as an effective mediator of communication between mechanical events occurring at different locations along the membrane[18, 17]. However, the specific details of the propagation of membrane tension appear to be system specific [15, 18, 10, 19]. For example, these views were recently challenged by the observation that membrane tension does not propagate in tissue culture cells, suggesting that the increased tension gradient remains locally confined to regions around the neck of the tether and does not relax for many minutes [15]. Likewise, in bacterial cells, the tension generated during tether pulling did not relax, indicating that the tension of the resting membrane is globally regulated and close to the lytic tension [20]. However, in axons of cultured neurons[18, 19], tension was shown to propagate many tens of micrometers.

This raises questions about how tension propagation is regulated along the membrane and what is the corresponding cellular adaptation mechanism to an external force. To provide insight into these questions, we established optical tweezer-based membrane nanorheology with a single, time-shared laser source[21, 22, 23] to study membrane tension propagation mechanics between a pair of lipid nanotubes extruded from a single axon. We used the well-studied touch receptor neurons (TRNs) of *Caenorhabditis elegans* as a biophysical model for tension propagation in axons. This model system has gained interest in the mechanophysiological regulation of animal behavior[24, 25, 26]. These neurons are important for the animal’s response to external forces. Although the exact mechanism of mechanotransduction is still not defined, genes affecting mechanoresponses are known and encode proteins of the extracellular matrix[27], microtubules[28], integral and peripheral membrane proteins[29, 30] and the cortical cytoskeleton[31], but also the membrane itself[32]. The spectrin cytoskeleton in particular has received ample attention and is known to organize associated membrane proteins into periodic clusters[33, 34], and is under constitutive mechanical tension[31]. While the role of the spectrin’s mechanical properties in mechanosensation is well known in *C. elegans*[31, 26], *Drosophila*[35, 36], chicken[37] and mouse[38, 39], the consequences of spectrin integrity on the propagation of membrane tension gradients is not understood. Likewise, stomatin homologs such as MEC-2 and STOML3, which contribute to mechanotransduction in the sensory neurons of *C. elegans* and mice [29, 40], bind cholesterol [41] and modulate membrane stiffness[40]. Strikingly, MEC-2 forms discrete condensates along the axonal membrane in vivo, arranged at regular intervals of 2–4 µm[28]. However, the functional implications of this periodic organization remain unclear.

To understand the link between the organization of these cellular elements and tension propagation, we combined our experimental results with a theoretical model proposed by [18, 15], in which tension propagation is associated with membrane lipid flow, but hindered by cytoskeleton-bound transmembrane proteins. The basic principle of our model is the same as those proposed before [18, 15], based on Brinkman flow, which extends the Stokes equation by adding a Darcy-like resistance term, allowing it to model viscous flow through porous media where both shear stress and permeability effects are significant [42]. Using this formalism, we conducted three-dimensional simulations using finite-element methods to compare the effect of membrane elasticity, viscosity, and permeability on tension propagation.

Our results suggest that membrane tension is locally confined and, accordingly, we reveal that tension propagation into the axon is limited by obstacles. Because we find that the extent of propagation differs between proprioceptive and touch mechanoreceptors that respond to different modalities, we propose that propagation speed and magnitude is an intrinsic property of the cell type and under tight genetic control.

### 1.1 Results

#### The active trap

To study how cell membrane mechanics enables tension propagation along the plasma membrane, we began our investigation with the classical tether extrusion experiment[43]. Similar to our previous approach with mechanosensitive proprioceptors[10], we used an optically trapped microsphere as a force probe and formed a contact between the probe and the membrane of isolated touch receptor neurons of *C. elegans* (Fig. 1a). Upon successful nonspecific attachment and withdrawal of the microsphere from the neurite, a membrane tether formed that required a constant force to maintain its position, reflecting the underlying baseline (or resting) membrane tension. A rapid retraction of the microsphere leads to an increase in force on the trapped bead, indicating the extension of the lipid nanotube (Extended Data Fig. 1a). After reaching a predefined distance from the neurite, we stopped the movement of the trap and held the probe at a fixed position. Immediately, the tether force rapidly decayed to a static value, which was different from the baseline value on the timescale of our experiment and independent of the initial extrusion velocity and, therefore, independent of membrane viscosity (Extended Data Fig. 1a), similar to what was observed before[43]. As noted previously, a higher retraction velocity caused an increased force on the microsphere[10], due to the finite viscosity and the resulting resistance of the membrane to flow into the tether (Extended Data Fig. 1b and [44, 45]). This viscosity-dominated peak increased with velocity up to 120 pN and was suggestive of the maximum tension difference of more than 1 mN/m generated during pull-out (Extended Data Fig. 1b,c). As the rapid increase in tension during the tether pullout is not due to fluid drag of the probe with the surrounding medium (see Methods), we reasoned that limited lipid flow into the tether caused this striking tension difference.

**Fig. 1.**
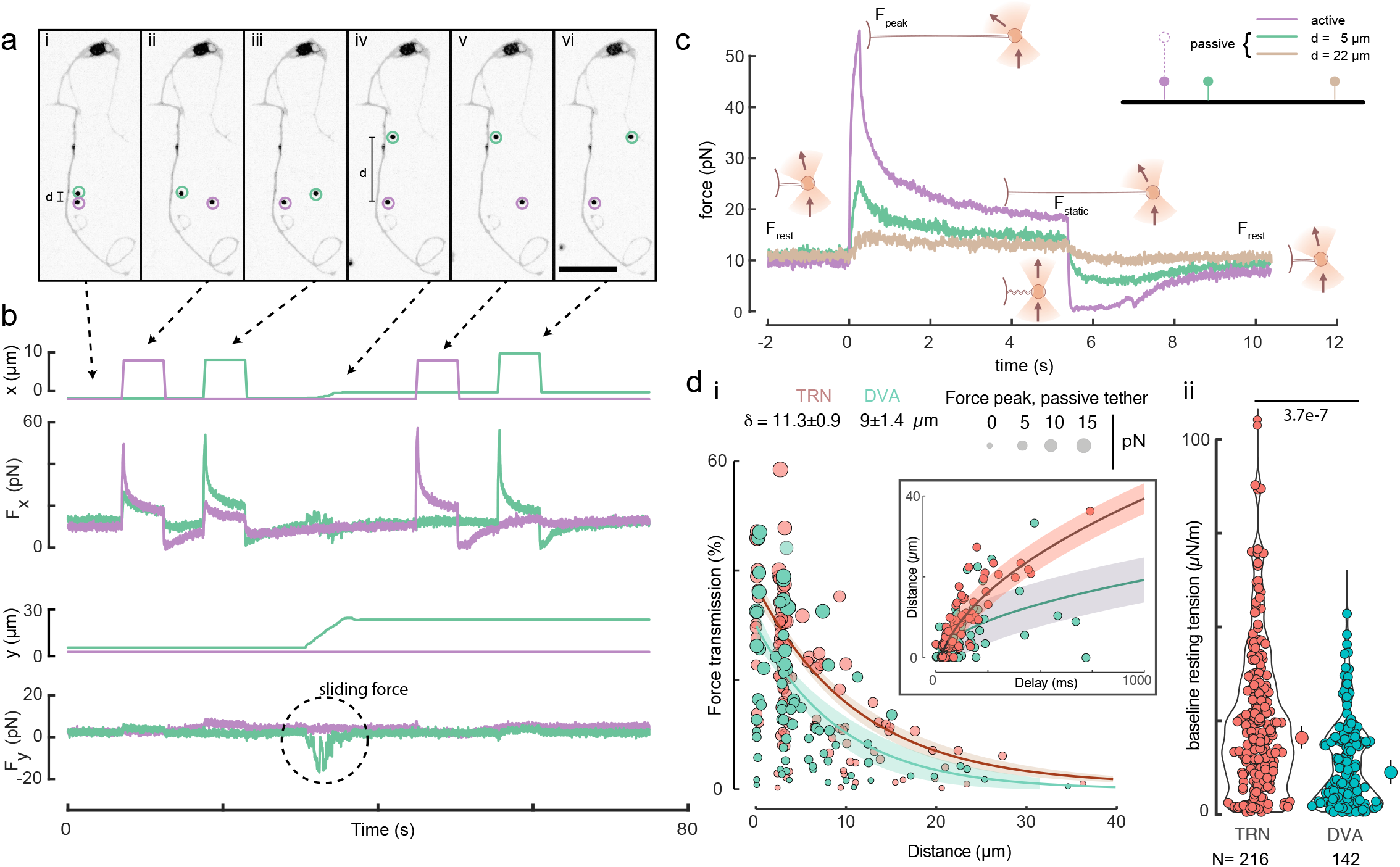
Adual trap assay to measure membrane tension propagation. **a**, Representative image of an isolated TRN with two membrane tethers extruded by two optically-trapped microspheres (purple and green circles for active and passive traps, respectively). Scalebar: 20 µm. (**i**) The two beads are connected to the axonal membrane through the extrusion of a lipid nanotube. (**ii, iii**) Both beads are sequentially pulled at 40 µm/s to a length of 10 µm (**ii**), while the resting, passive trap records changes in membrane tension at a distance *d*. (**iv-vi**) *d* is increased to record tension propagation under varying separation. **b**, X-(top) and Y-(bottom) component of the trap trajectory and force. The sequence events depicted in (a) are indicated with black arrows. The sliding friction (*F*_*y*_ component) between events iii and iv is indicated with a dashed circle. **c**, Simultaneous force measurement during trapezoidal pulling routines for the *active* trap and *passive* trap placed at *d* = 5µm (event ii in a, b) and *d* = 22 µm (event v in a,b). While a small increase in membrane tension in the *passive* trap is detected when this is placed close to the active trap, membrane tension at *d* = 22 µm shows no force peak but a slow, second-scale tension increase. The force drop *F*_*x*_ *→* 0 at *t* = 5 indicates tether buckling upon the fast approach of the trap towards the axon. Trap velocity is faster than the ability of the axon to absorb the lipidic material from the tether. Afterwards, tension homeostasis induces tension recovery. Schematic inside the plot graphically illustrate the events and the deflection of the laser due to bead displacement from the trapping center. Thick arrows indicate direction of the laser. **d**, (i) Change in peak membrane tension measured at the *passive* lipid nanotube for increasing distances normalized to the *active*, pulling site, calculated with Eq. 6. The size of the points corresponds to the absolute transmitted force at the passive tether. Solid line is the exponential fit with the error band indicating the 95% confidence interval of the fit; *δ* indicates the characteristic length-scales derived from the exponential fit; p-value derived from comparing a linear regression on pooled data with interaction term (see Methods). Red points indicate values derived from TRNs, green points from DVA. Inset shows the tension propagation velocity for the two cell types. Solid line indicates best fit of *t*^0.5^ to the data with the 95% confidence interval. (ii) Resting plasma membrane tension of TRN vs. DVA cultured neurons. The median of the distribution is shown as a circle to the right of the violin with the vertical bar indicating the 95% confidence interval. The p-value derived from a two-sided Kolmogoroff-Smirnoff test. N= number of tether extrusion events.

When the microsphere is moved again towards the neurite such that the membrane tether is released back into it, the force drops below its resting value indicating negative tension differentials (Fig. 1b,c). However, the baseline, resting plasma membrane tension recovered within 2-3 s (*F*_*rest*_ ≈ 6-9 pN), indicating a mechanism that maintains tension homeostasis. The resting force *F*_*rest*_ is related to the membrane tension *σ*_*rest*_ through the bending rigidity of the membrane *κ* according to

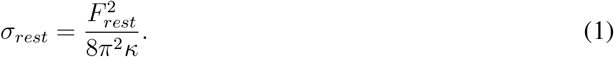

Thus, using this simple relation, we can approximate the tension in the membrane by measuring the static forces on the bead in the trap[43, 46].

At high extrusion velocities, we observed spherical instabilities forming within the extruded membrane tube. These instabilities originated near the tether neck and propagated toward the microsphere (Video 1, Extended Data Fig. 1 d). Similar dynamics are frequently observed in mechanical instabilities [47]. This instability traveled along the tether, indicative for a gradient in mechanical tension driven transport. Interestingly and in contrast to previous observations[18], the instability never visually propagated to the neuronal process, indicating that the increase in membrane tension was locally confined and hardly propagated into the neurite (Extended Data Fig. 1d). From these results, we concluded that there exists a strong tension difference inside the tether, compared to the resting tension of the axonal membrane far from the tether, which does not, or only incompletely, propagate into the neurite. This difference gives rise to a tension gradient over the length of the tether, which relaxes as lipid flows from the membrane reservoir into the tether. Under constant tether length, radial expansion driven by lipid transport reflects the classical inverse scaling between tether force and radius[48].

#### The active and the passive trap

To gain insight into the mechanism of tension propagation in neurons, we directly measured the extent of tension propagation along the axon to a point perturbation. To do so, we established an optical trapping experiment to simultaneously extrude two membrane tethers with a single, time-shared laser source (Fig. 1a, Video 2, Extended Data Fig. 2a, Supplementary Text 1). Similarly to our previous time-shared optical tweezers microrheology scheme (TimSOM, [23]), we split a single laser source at 25 kHz to generate two traps that can be moved independently in 2D (Fig. 1a, Extended Data Fig. 2, SI Text 1). Due to the asynchronous operation of the two traps in the time-shared configuration, we assessed the impact of temporal multiplexing on force measurements. We simulated the bead’s response to an external tether force within a periodically switching optical trap, which predicted a systematic underestimate, independent of the measured force (Extended Data Fig. 2b, SI Text). This prediction was experimentally validated by comparing tether extrusion events conducted in a continuously active trap versus a time-shared trap. The measured forces in the time-shared configuration were consistently underestimated by a factor of 0.63, which we applied as a correction factor throughout our analysis. Using these two effectively simultaneous optical traps, we extruded a single membrane tether with each of the two traps that were placed in close proximity to each other (Fig. 1 a). Then, we actively pulled on the tether in trap 1, while measuring the response in the static tether, held with trap 2 (Fig. 1b). We alternated this procedure for trap 1 and trap 2 and repeated the extrusion protocol for dual tether distances (DTD) of up to 40 µm (Fig. 1c, Video 2). As expected, the baseline tension measured in the active trap and in the passive trap is indistinguishable, demonstrating that the tension of the unperturbed membrane is homogeneous throughout the neurite (Extended Data Fig. 3a).

In addition to the active trap described above, the passive trap also displayed rich behavior. At close separations from the active trap (directly opposite to the active trap), we observed a strong reduction in the propagated peak force (Fig. 1c). Only ≈40-50% of the force of the active trap reached the passive trap, even at distances less than 1 µm. The force propagation velocity from the active to the passive tether was very rapid and, in contrast to previous work[15], exceeded 150 µm/s. However, the time required to reach the passive trap increased non-linearly with distance (inset of Fig. 1d), suggesting that the propagation velocity of the mechanical pulse decreases as it travels along the neurite (Extended Data Fig. 3b-e; Methods). We also observed considerable variability in the propagation length scale (Fig. 1d) and velocity (Extended Data Fig. 3e), which may reflect the involvement of distinct propagation mechanisms—such as transmission through the membrane, the cytoskeleton, or the viscoelastic cytoplasm.

With increasing distance from the active tether, the peak of the passive trap decreased substantially (Fig. 1 d) until it seemed to disappear. At *>* 20 µm tether separation, the peak was completely buffered. Instead, we observed a slow and steady (≈second scale) increase to a plateau value (brown curve in Fig. 1c), suggesting that the membrane behaves as a low-pass filter for mechanical signal transmission only the slow components get propagated up to 40 µm. This plateau force does not significantly depend on the distance to the active tether (Extended Data Fig. 3f). To calculate the extent of force propagation, we plotted the ratio of the peak force in the passive tether to the peak force in the active tether as a function of the distance *x* between the two tethers (Figs. 1d, Extended Data Fig. 3) and fitted an exponential decay function exp(*x*) = *A* exp(−*x/δ*) using weighted nonlinear least squares to the absolute peak force of the passive tether (see Methods for details), where *δ* is the characteristic length scale of the tension propagation and *A* as the amplitude of the ratio. The weighted exponential (Akaike Information Criterium, AIC: 854) fitted the data better than a Gaussian (diffusion) kernel (AIC: 860) or a hyperbolic function. Thus, we used an exponential function to quantify tension propagation distances and mechanistically dissect the governing mechanical properties.

We then sought to understand whether tension propagation is different in mechanoreceptor neurons with different functions and turned to DVA proprioceptors. The DVA neuron differs from TRNs in that it contains a single axon, which senses local changes in body bending to coordinate body curvature[49, 10]. We previously showed that higher tension gradients during tether extrusion transiently suppress calcium signals in DVA through the two-pore potassium channel K2P TWK-16[10]. Conversely, tether relaxation and the resulting negative membrane tension gradients initiate local calcium transients that propagate along cultured neurites via the mechanosensitive NOMPC TRP-4 channel [10]. This model, in which a local reduction in membrane tension after the relaxation of the tether (e.g., negative tension gradient) activates NOMPC TRP-4 under mechanical compression [50, 51] and positive tension—and membrane stretch—opens K2P channels [52], provides a coherent framework for bidirectional mechanosensory tuning[53]. We thus speculated that mechanical stresses compartmentalize the axon into dynamic active zones. One requirement of this model is that the spread of mechanical information is limited within the axonal membrane. By applying our dual tether assay, we found that the resting tether force is lower compared to TRNs (Fig. 1d ii; p = 3.7 × 10^−^7, two-sided KS-test) and that the tension remained more confined in DVA and did not propagate far (Fig. 1d i) when compared to the measurements in TRNs (Fig. 1d, 11.5 vs 9 µm). Intriguingly, we found that the propagation velocity was lower in DVA than in TRNs (Fig. 1d), indicating slower transport of mechanical information along the membranes of proprioceptive neurons. Touch receptor neurons are known to be exquisitely responsive to high-frequency stimulation above 10Hz [54, 55, 56], whereas DVA also responds to slower (< 1Hz), proprioceptive cues such as undulatory body motion[49, 10]. Thus, the transport mechanics may impose a lower bound on the frequency response, helping neurons selectively respond to different stimulation velocities. Together, this indicates that membrane tension propagated further and faster in TRNs. More generally, our data show that the extent and speed of membrane tension propagation is cell-type specific, which motivated us to dissect the molecular and mechanical mechanism of tension propagation in neurons.

#### Membrane tension propagation is restricted by membrane obstacles

Following recent attempts to describe tension propagation quantitatively[15], we assumed that membrane lipid flow transmits local changes in tension and that cytoskeleton-bound transmembrane proteins, such as cell adhesion molecules, ion channels, and membrane-cortex cross-linkers, act as obstacles impeding membrane flow (see Section 4.4 in the Methods). These dynamics are described by

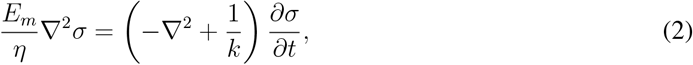

where the membrane tension, *σ*, represents the force per unit length acting on a membrane cross-section. This force counteracts membrane stretching or compression. The effective area expansion modulus of the cell membrane, *E*_*m*_, is a material property of the membrane, and it specifies the relationship between the force applied to the membrane and the resulting change in its surface area. *η*, the two-dimensional membrane viscosity, resists the flow of lipids and other molecules inside the membrane. The Darcy permeability of the obstacles, *k*, measures the ease with which tension propagates through the membrane (Fig. **??**a). We used a cylinder as an idealized geometry for a segment of the axon, and the tethers were represented by attached thinner cylinders (Fig. **??**a). As in [15, 18], we defined the permeability *k* = *a*^2^*f* (*ϕ*) by the obstacle area *a* and a function of the obstacle density *ϕ, f* (*ϕ*) = − (1 + ln(*ϕ*))*/*(8*ϕ*). This function was derived from the calculation of the mean force required to drag one particle with mean velocity through the background of immobile particles, see [18] for details. For typical values of *ϕ* < 0.3 [18], *f* is a decreasing function of obstacle density. Thus, increased permeability results from a reduced number of obstacles covering the same area, as well as an increase in obstacle size, which can occur due to molecule clustering (Fig. **??**b).

From Eq. 2, we observed that tension propagation is governed by the parameters *E*_*m*_, *η*, and *k*. Hence, we examined the effect of changing these parameters in the tension propagation using finite element simulations in a cylindrical geometry (see Materials and Methods for details). To quantify the extent of tension propagation, we fitted the same exponential decay model from the experimental data (*A* exp(−*x/δ*)) to the simulation data. However, there was a systematic underestimation of the exponential fit function for large distances between tethers in the simulations. Adding an offset parameter *C* to the exponential decay (i.e., *A* exp(−*x/δ*) + *C*) resulted in better fittings (for *k* = 0.03 µm^2^, the AIC without offset is 42.99 and the AIC with offset is 8.24) and reflects the propagated tension at long distances. Additionally, this offset in the simulations accounts for processes not represented in the model, such as material sources and sinks. When permeability *k* is high, there are fewer and/or smaller obstacles in the membrane, allowing it to flow more freely. Consequently, the propagation of tension increases as the permeability increases (*k*, Fig. **??**c, Extended Data Fig. 4a). Because our model also incorporates membrane elasticity, we asked to what extent *E*_*m*_ affects tension propagation. We found that the more elastic the membrane, the greater the tension at the tether, which increases the extent of tension propagation along the axon (Extended Data Fig. 4b). Lastly, we hypothesized that a higher viscosity hinders the lipid flow of the membrane and, hence, localizes the tension propagation to a smaller region. Indeed, when the viscosity *η* increases, we observed a decrease in tension propagation (Extended Data Fig. 4c). Together, our simulations suggest a striking dependence of tension propagation on membrane in-plane elasticity, viscosity, and obstacle density.

Recent studies have shown that ion channels are organized in a periodic arrangement of obstacles along the axon [34] and dendrites [57] of neurons from various organisms, including *C. elegans* [26]. Our initial model assumes that obstacles are randomly and homogeneously placed in neurites, which may not reflect the physiological organization in neurons. The periodic arrangement is governed by the actin-spectrin cytoskeleton, in which the spectrin tetramers expand ≈190 nm and connect adjacent actin rings (Fig. **??**d). We introduced this periodic arrangement along the axon by assuming that although ion channels and transmembrane proteins are present along the axonal membrane, they assemble into different clusters, based on experimental observations [34]. Moreover, we assumed that 2/3 of the 190 nm periodic interval is covered by channels and 1/3 by smaller obstacles attached to the actin rings (Fig. **??**e, [34]). We proposed that permeability *k* in the zone related to actin rings is higher *k*_+_ = 1.5*k* than in the zones covered by channels *k*_−_ = (3*k −k*_+_)*/*2 due to the larger projected area of often multimeric ion channels (e.g Piezo2, [58]) compared to single-pass transmembrane proteins or G-protein coupled receptors (Fig. **??**e). We varied the parameters and observed that tension propagated less in the periodic control case, i.e. using *k* from the homogeneous control case to calculate *k*_+_ and *k*_−_, than in the homogeneous control case (Fig. **??**f). In all cases, the ratio of the passive to the active tension peak is higher compared to the random obstacle arrangement. This indicates that a periodic arrangement localized tension and leads to less propagation along the axon. Indeed, our parameter sweep confirmed this assessment even though the parameters show the same tendency as in the homogeneous, randomized organization (Extended Data Fig. 4 b, d, and, e). A consequence of the periodic arrangement is that tension decays faster with distance, leading to a more confined mechanical tension propagation. Note that the trend is clear for all the fitting results of the length scale and that the random and periodic arrangements are statistically significant for *k* = 0.02, 0.03, 0.04 µm^2^, and some values of *η* and *E*_*m*_ (Extended Data Fig. 4a, f, g). Because the simulations are deterministic, the shorter tension propagation in the periodic case is a causal consequence of the change in spatial organization of the obstacles and is highly relevant for the functional outcome of the mechanical properties. Together, our simulations show that periodic obstacles can curb the extent of tension propagation along axonal membranes.

We reasoned that, due to the complex nature of membrane–cortex interactions, the model parameters cannot be directly attributed to specific membrane or cytoskeletal components. Instead, tension propagation appears to be governed by multiple synergistically acting parameters. For example, changing the actin organization and membrane cortex adhesion may disrupt obstacle density but also change the viscosity [3]. To better understand how these parameters jointly influence tension propagation, we performed simulations by varying two parameters simultaneously. Fig. **??** g-i shows that the tension can propagate further when the parameters are changed simultaneously. For example, in Fig. **??**g, decreasing *η*, results in larger tension propagation for any value of *E*_*m*_, but if *E*_*m*_ is increased, the tension propagates further. This suggests a cooperative effect of both parameters, *η* and *E*_*m*_, on tension propagation. Moreover, the degree of tension propagation depends on the ratio *η/E*_*m*_. Note that in Eq. 2, this ratio gives the time scale of the dynamics. In summary, we detected a stronger synergistic relationship between the parameters *k* and *η* and that the periodic arrangement of the obstacles affected the propagation of tension in silico. We next investigated the role of these properties systematically in experiments.

#### Lipid saturation has little influence on tension propagation

Through AFM-based tether pulling experiments [45] from the cell body of touch receptor neurons, we have previously shown that lipid saturation modifies the viscous properties of the membrane but not the baseline resting tension [32]. Therefore, we asked whether lipid saturation and osmotic effects influence tension propagation along the neurite. *fat-3* mutant neurons with defects in polyunsaturated fatty acid (PUFA) synthesis [59, 32] did not have a discernible effect (Fig. 2 a, b). However, because our tissue culture medium contained 10% fetal bovine serum, the effect of *fat-3* may be partially confounded by the PUFAs in the medium[60]. Recently, intracellular pressure was shown to influence the rate at which tension propagated along the membrane in HEK293T cell cultures [61]. We also observed that the application of a hypoosmotic shock significantly increased the tether force (p=0.007). However, we detected no significant difference in tension propagation compared to wildtype (Fig. 2c; *δ*= 11.3 vs 11.5µm). Together, these data show that tension propagation is surprisingly robust to subtle changes in the composition of the lipid bilayer and increases with intracellular osmotic pressure. Furthermore, it suggests that tension propagation may be dominated by obstacles in the membrane and cytoskeleton.

**Fig. 2.**
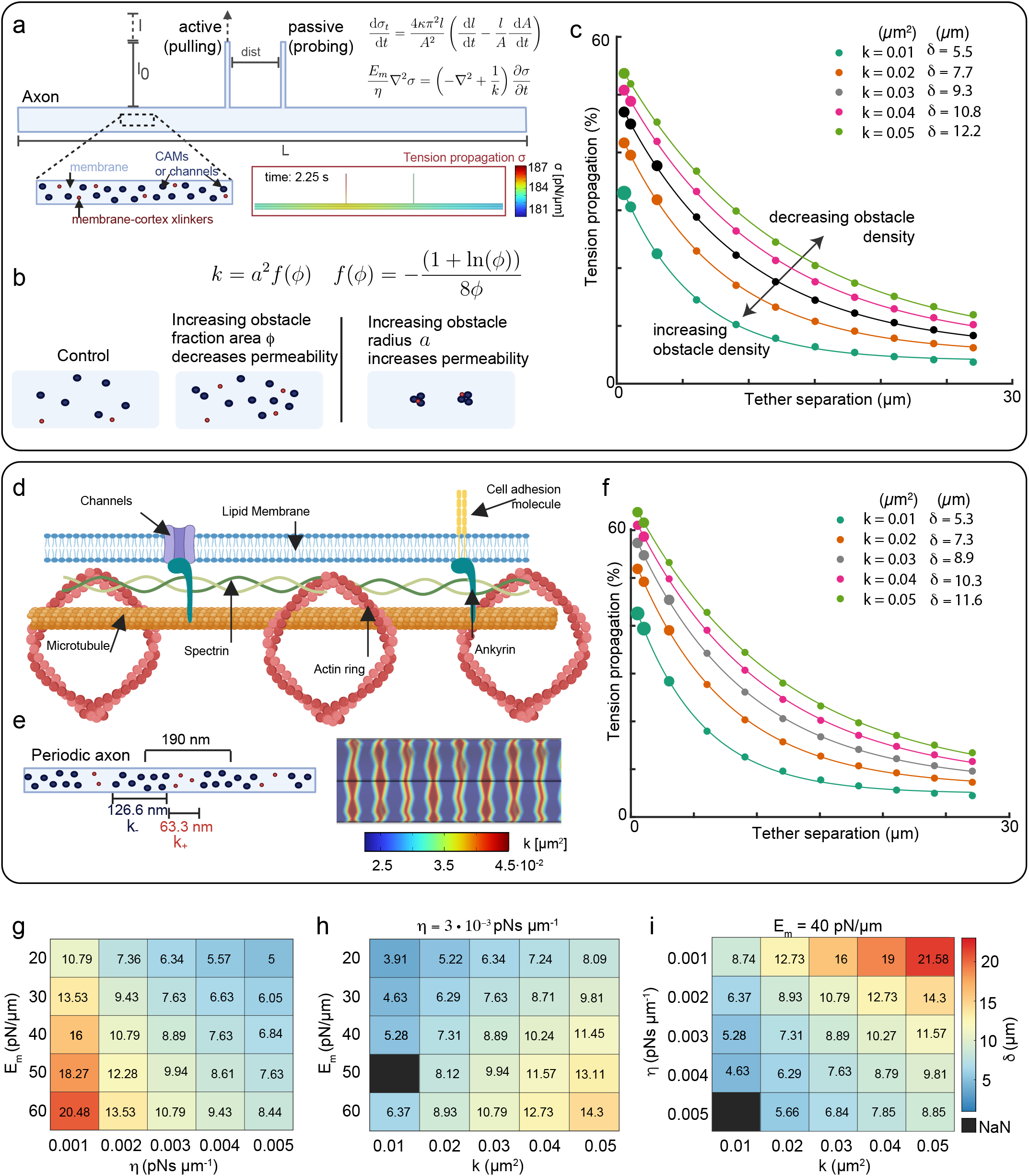
Theoretical model for tension propagation. **a**, Schematic representation of the modeled axon: tension *σ* generated by pulling the active tether *σ*_*t*_ propagates along the axon. Left inset shows a random distribution of membrane obstacles. The right inset is a snapshot of the simulation at time 2.25 s. The distance between tethers is 3 µm and the colorbar represents the tension *σ*. CAM, cell adhesion molecules. **b**, Top view of the membrane (a), showing possible ways to change the permeability *k*. **c**, Change in membrane tension measured in the *passive* lipid nanotube for increasing distances normalized to the *active*, pulling site, with different permeabilities due to randomly distributed obstacles. The size of the circle corresponds to the amount of tension in the passive tether in the simulation (small 0 *< σ ≤*3pN/µm, medium 3 *< σ ≤*6pN/µm, and big 6 *< σ ≤*9pN/µm). Black represents the control condition. **d**, Lateral view of the axon showing its underlying molecular structure. Figure created in BioRender. **e**, Periodic distribution of obstacles along the axon and the corresponding values of *k*_+_ and *k*_−_. The image on the right shows the color-coded values of *k*_+_ (red) and *k*_−_ (blue) along the axon in the simulation. Note that the values are interpolated and follow the underlying triangular mesh. **f**, Change in membrane tension using periodic configuration for different parameters. Note that *k* corresponds to the mean value of the permeability along the axon that takes *k*_+_ = 1.5*k* and *k*_−_ = (3*k −k*_+_)*/*2. The gray color corresponds to the control condition. **g-i**, Two dimensional phase diagram of tension propagation in presence of periodically arranged obstacles with synergistically varying (g) *E*_*m*_ and *η*, (h) *E*_*m*_ and *k* and, (i) *k* and *η*. The corresponding parameter values are denoted on the vertical and horizontal axes. Colors represent the extent of tension propagation *δ*. The exact values of *δ* are added to the heatmap. NaN (black squares) represent conditions for which simulations at certain displacements did not converge.

#### The scale of tension propagation is modulated by the cytoskeleton

In our simulations, we treated the membrane and the underlying cytoskeleton as a composite material and assumed that changes in the cytoskeleton affect membrane permeability. This is because transmembrane proteins, which connect to the actin cytoskeleton, can act as obstacles to the flow of lipids in the membrane, as suggested in previous work [62]. Therefore, we measured tension propagation and static tension in four different treatments with Latrunculin A (LatA), a pharmacological agent that sequesters actin monomers and is known to interfere with actin polymerization [63], and also Cytochalasin D (CytoD), a drug known to depolymerize dynamic actin filaments [64]. At low concentrations of LatA (delivered at 0.5-1 µM acutely, 30 min incubation time), we did not observe an effect on membrane tension (Fig. 3a) or on tension propagation (Fig. 3b, Extended Data Figs. 5 and 6). This is consistent with earlier studies showing that low concentrations of LatA does not disrupt stable actin filaments and does not interfere neither with membrane tension extruded from the neuronal cell bodies [31] nor with actin ring disassembly in *Drosophila* neurons [65]. Notably, acute application of 10 µM LatA resulted in an unexpected increase in the baseline resting membrane tension (Fig. 3a) that was accompanied by a significant reduction in tension propagation (see Table 1). This observation was remarkably different upon acute delivery of 2 µM CytoD, which effectively promoted tension propagation (Fig. 3b), suggesting that acute actin filament capping (e.g, CytoD) and monomer sequestration (e.g. LatA) affected membrane permeability differently. However, after chronic incubation of neurons with 1 µM LatA or 2 µM CytoD overnight, we observed a significant and robust reduction in static membrane tension (LatA: p=210^−7^; CytoD, p<110^−16^; Fig. 3a). The propagation of membrane tension was also differently affected by different drug concentrations. We found that chronic application of high concentrations of LatA and CytoD resulted in very limited tension propagation (ctrl vs. LatA vs. CytoD; 11.3 vs 8.1 vs 8.1 µm, Fig. 3b). Overall, this is consistent with our previous observations and other reports, which suggest that neuronal responses to actin disruption are drug- and concentration-dependent, with greater sensitivity to cytochalasin D (CytoD) compared to acute treatments with latrunculin A (LatA) [31, 65].

**Fig. 3.**
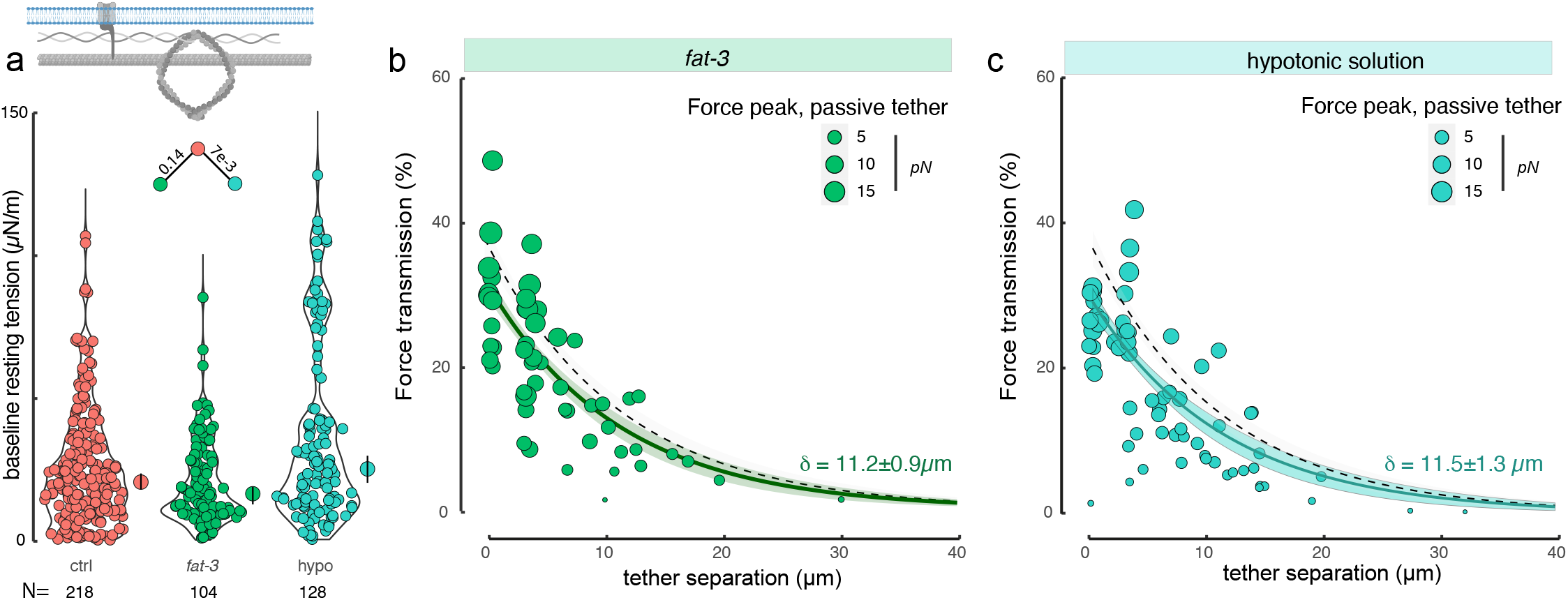
Hypotonic shock but not lipid saturation modulates tether forces extruded from neurites. **a**, Violin plot of the baseline, resting membrane tension during the membrane extrusion for control cells, *fat-3* mutants, that perturb lipid composition, and on cells treated with hypotonic shock. Median shown as circle with vertical bar indicating the 95% confidence interval next to the violin distribution. N= number of measurements. Top schematic highlights membrane in celeste. p-values derived from a two-sided Kolmogoroff-Smirnoff test above the color-coded combinations. **b, c**, Tension propagation in the membrane of cells mutant for (b) *fat-3* and (c) cells treated with a hypotonic solution. Solid lines indicate the fit to the an exponential decay, shaded area indicating the 95% confidence interval of the fit as described in the methods, dotted black line indicates the fit derived from control data (Fig. 1d). Size of the dots corresponds to the absolute force transmitted to the passive tether. Fit parameter *δ* indicates the characteristic length scale.

We reasoned that different concentrations of latrunculin and cytochalasin perturb different populations of actin filaments, as previously observed in MDCK cells [66]. This could be understood in transient decoupling of the cortex membrane (increase in *k*) with low concentrations of drugs, followed by a decrease in membrane elasticity (decrease in *E*_*m*_) in chronic drug application. To understand how propagation is subject to the mechanical properties of the membrane-cytoskeleton composite, we turned to our model and simulated different conditions of *E*_*m*_ and *k*. In fact, a decrease in membrane elasticity leads to a decrease in propagated tension, where an increase in *k* led to increased propagation of tension (Extended Data Fig. 5 and Fig. 3c).

Having established the role of actin in axonal membrane mechanics, we next investigated whether microtubules modulate tension propagation along the membrane. Microtubules are a hallmark of neurons and are known to interact with mechanosensitive ion channels[67, 68] and membrane-associated proteins[69], making them strong candidates for influencing membrane tension.

In *C. elegans* TRNs, the microtubule cytoskeleton includes large, crosslinked bundles of specialized 15-protofilament microtubules, which are highly acetylated[70, 71], in addition to the conventional 11protofilament type. These bundles are anchored to the plasma membrane via a spectrin/ankyrin-based complex[72].

To test their role, we pulled membrane tethers from TRNs of *mec-12(e1607)* mutants, which lack the acetylated 15-pf microtubules but retain conventional ones[73]. These mutants showed significantly increased baseline membrane tension (p = 0.002). One possibility is that disrupting the acetylated bundle activates cortical contractility via Rho signaling, as seen in cultured cells[74]. However, a more likely explanation is that 11-pf microtubules compensate by altering membrane attachment. Supporting this, treating *mec-12* mutants with nocodazole reversed the elevated membrane tension (Fig.3 d).

Interestingly, *mec-12* mutants also exhibited reduced tension propagation compared to controls (10 vs. 12.3µm), and nocodazole treatment led to an even greater reduction (p = 0.01; Fig. 3e,f). These results demonstrate that microtubules support membrane tension propagation in axons and suggest that their loss affects propagation through at least two distinct mechanisms: changes in cytoskeletal rigidity and disruption of membrane anchoring.

#### The periodicity of the actin/spectrin cytoskeleton sets limits on tension propagation

It is well-established that dynamic barriers on the cytoplasmic leaflet hinder lateral membrane movement[75]. The actin-spectrin cytoskeleton is important in that it coordinates many transmembrane proteins (ion channels, adhesion receptors, dystroglycans, band 4 proteins) and also binds to the membrane itself [76, 34, 77], effectively compartmentalizing the membrane[62]. Thus, it is plausible that the periodic spectrin cytoskeleton establishes a barrier that impedes free tension propagation by establishing periodic obstacles. Indeed, when we simulated periodically arranged obstacles, rather than randomly distributed clusters, the overall tension was higher but attenuated over short distances and therefore propagated less (Fig. **??**f). To better understand how obstacle organization influences tension propagation, we simulated tension dynamics increasing the value of *k*_+_ to 2.6*k*, which in turn decreases the value of *k*_−_ = (3*k −k*_+_)*/*2. These changes accentuate the difference between the permeabilities and hence, represent a more predominant periodic barrier. We compared the outcomes between randomly distributed obstacles and the new setup of periodically arranged ones (Fig. 4b). These simulations revealed that periodic cytoskeletal arrangements enhance tension propagation over short distances. However, at intermediate distances, which depend on specific model parameters such as permeability (*k*) and membrane elasticity (*E*_*m*_), random arrangements favored greater tension spread (Fig.4b, inset). In In other words, loss of periodicity leads to an increase in tension propagation, especially at large distances.

**Fig. 4.**
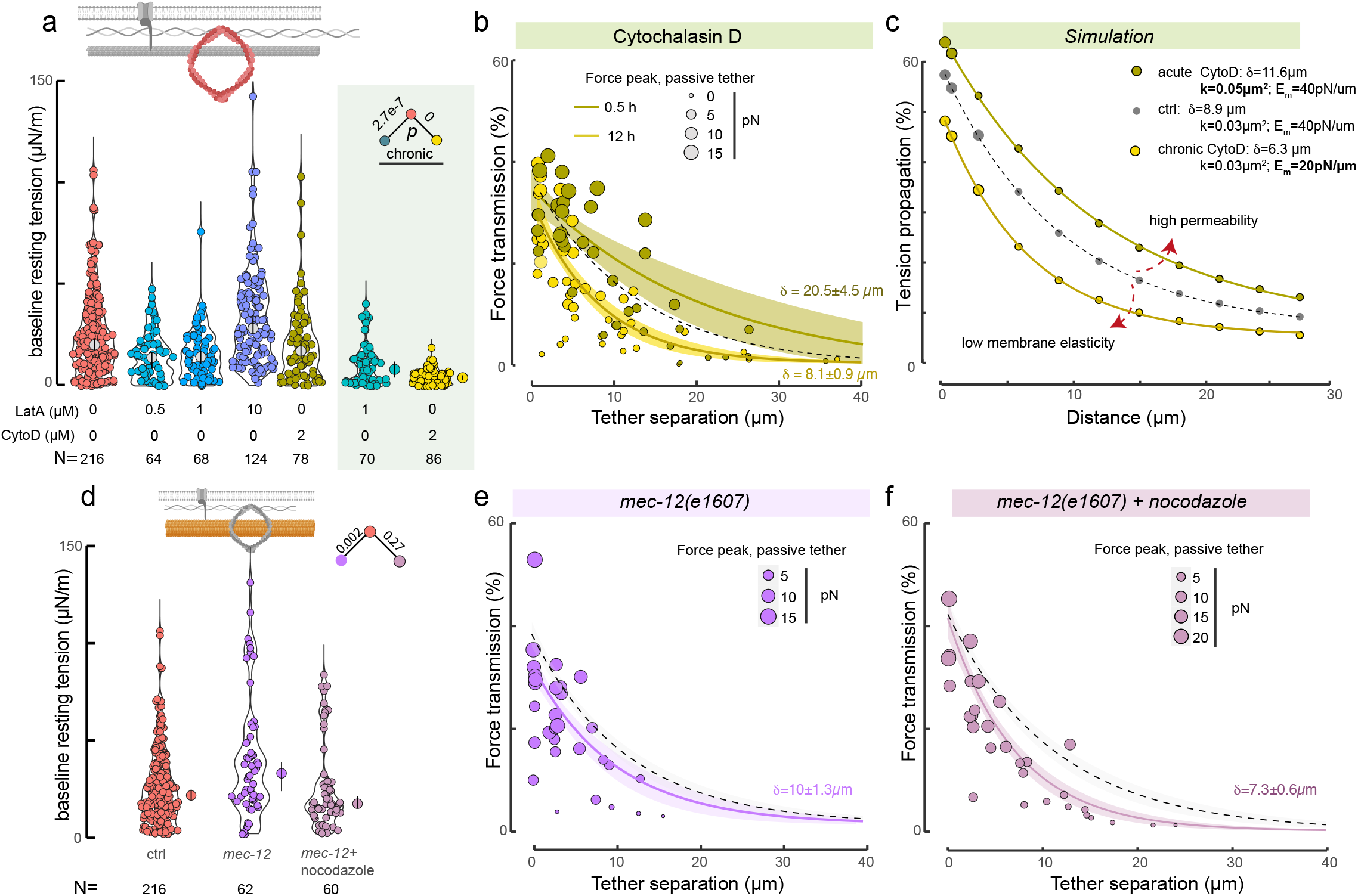
Tension propagation is sensitive to F-actin and microtubule network integrity. **a**, Influence of the F-actin cytoskeleton on baseline, resting membrane tension in cells treated with different concentrations of cytochalasin and latrunculin to perturb the F-actin cytoskeleton. Chronic incubation is performed over night (12h). **b**, Tension propagation in cells after acute (green) and chronic (red) incubation with cytochalasin D. **c**, Simulation of tension propagation with reduced *E*_*m*_ and increased *k* compared to control, describing chronic and acute actin depolymerization. Parameters used for simulation are shown in the figure. The simulation corresponds to the periodic case. Hence, the value *k* is used to calculate *k*_+_ and *k*_−_. See also Extended Data Fig. 4. **d-f**, Influence of the microtubule cytoskeleton on (d) static membrane tension and tension propagation in the (e) *mec-12* mutant and cell treated with nocodazole. For panels (a) and (d): The top schematic highlights different cytoskeleton elements of interest. The median is indicated as a circle with a vertical bar indicating the confidence interval 95%. N = number of measurements. p-values above the color-coded combinations derived from a two-sided Kolmogoroff-Smirnoff test (see also Table 1). For panels displayed in (b), (e), and (f): solid lines indicate a fit to an exponential decay and the shaded area indicating the 95% confidence interval of the fit as described in the methods, dotted black line indicates the fit derived from control data (Fig. 1d). The size of the dots corresponds to the absolute force transmitted to the passive tether. Fit parameter *δ* indicates the characteristic length scale.

UNC-70 is the sole *β*-spectrin homolog in *C. elegans* neurons and forms a complex with *α*-spectrin, and can bind to both the actin cytoskeleton and the membrane with its pleckstrin homology domain [78]. In *C. elegans*, similar to the neuronal cytoskeleton in mammals, both components form a characteristic periodicity of 190 nm in vitro and in vivo [26, 34, 57]. This periodicity is more variable in intact animals[79] and completely lost in the absence of functional UNC-70, as visualized using *α*-spectrin as a proxy for the spectrin network organization (Fig. 4a and [26]). Similarly to model predictions (Fig. 4b), the propagation at short distances in the spectrin mutant is lower than in the wildtype, but propagates significantly further into the neurite of *unc-70* null mutants (Fig. 4c; p=0.02), suggesting that the loss of *β*-spectrin leads to discretely clustered domains of transmembrane proteins [80] and concomitant increase in permeability.

However, our model does not rule out other factors, such as transmembrane receptor endocytosis, which may further lead to an increase in tension propagation in the spectrin mutant [81]. Together, our results from both the experiments and modeling suggest that periodic membrane obstacles as well as components of the cytoskeleton locally restrict and limit the extent of membrane tension propagation. Our observations suggest that the distinct periodicity of the actin/spectrin network is a mediator of the organization of transmembrane and membrane-associated proteins and thus the propagation of tension in neurons.

#### Stomatin MEC-2 condensation at the membrane impedes tension propagation

Lastly, we tested whether obstacles in the membrane can indeed influence the propagation of membrane tension. We have previously shown that MEC-2 (*mec* for *mec*hanosensory), a stomatin homolog associated with stiffened membrane domains[40], forms biomolecular condensates proximal to the inner leaflet of the plasma membrane[29]. MEC-2 condensates exhibit a discrete, punctate distribution in both in vivo and in vitro settings, with a typical spacing of 2-4 µm and 1-2 µm (Fig. 5b inset), respectively. To understand whether condensation of MEC-2 on the plasma membrane influences tension propagation, we performed tether extrusion in *mec-2* knockout animals. The baseline tension of the resting membrane was lower than that of the control neurites, indicating that the MEC-2 stomatin may also influence tension propagation (Fig. 5 a). Interestingly, we also observed substantially larger tension propagation in the neurite of *mec-2* mutants (ctrl vs. *mec-2*; 11.3 vs. 18.5 µm). Thus, we asked whether obstacles like MEC-2 impede tension propagation in our model. Indeed, as shown above, our simulations predicted that membranes with a higher permeability would enhance tension propagation (Fig. 5c). Therefore, removing MEC-2, which binds to the inner leaflet of the plasma membrane through its cholesterol binding ability, reduces obstacle density and increases *k*, also increases tension propagation.

**Fig. 5.**
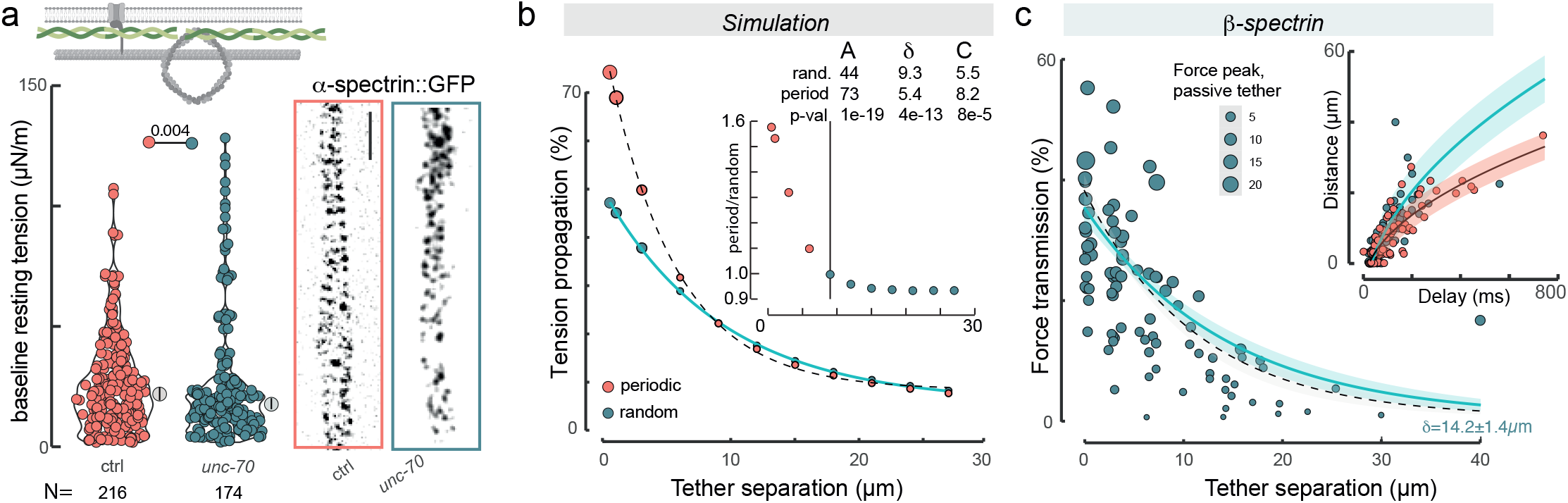
Periodicity of the spectrin cytoskeleton supports tension propagation over short distances. **a**, Influence of the spectrin cytoskeleton on static membrane tension. The median is indicated as a circle with a vertical bar indicating the confidence interval 95%. N = number of measurements. p-values above the color-coded combinations derived from a two-sided Kolmogoroff-Smirnoff test (see also Table 1). The images on the right show representative superresolution images of *α* -spectrin in wild-type and *unc-70* mutant neurons. Scalebar = 1 µm. **b**, Simulation of the tension propagation comparing cells with periodic (red circles) and random (blue circles) obstacle arrangement. Fit parameters of the black, dotted line (periodic) and the blue solid line are indicated in the lower left. The inset shows the ratio of the propagated tension in the periodic relative to the random case, showing that periodic arrangement favors tension propagation over short distances, while random arrangement favors long-distance propagation. The following values for permeabilities have been used: *k*_+_ = 2.6 *· k* (instead of *k*_+_ = 1.5 *· k*), and *k*_−_ = (3*k −k*_+_)*/*2). Thus, the value of *k*_+_ increases (0.078 from 0.045 µm^2^) and the value of *k*_−_ decreases (0.006 from 0.0225 µm^2^), with k = 0.03 µm^2^. **c**, Tension propagation in *unc-70* mutant cells. Inset: tether separation vs. pulse delay. The fit shown as solid line and confidence interval shown as a shaded band indicate pulse propagation velocity. Solid lines indicate a fit to an exponential decay as described in the methods and the error band indicating the 95% confidence interval of the fit, dotted black line indicates the fit derived from control data (Fig. 1d). The size of the dots corresponds to the absolute force transmitted to the passive tether. Fit parameter *δ* indicates the characteristic length scale.

**Fig. 6.**
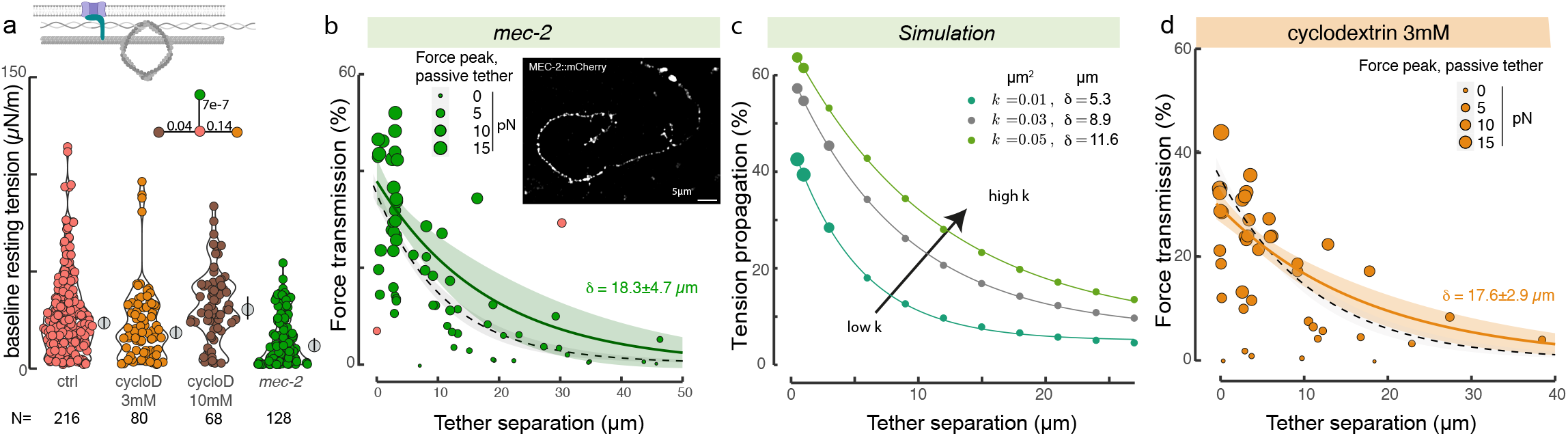
Membrane condensation of MEC-2 delimits tension propagation along the membrane. **a**, Baseline, resting tension during the membrane extrusion from cells mutant for *mec-2* or treated with increasing concentrations of methyl-*β*-Cyclodextrin (cycloD; 3 or 10mM, see Methods). Top schematic highlights channel-cytoskeleton crosslinkers in blue. Median is shown as circle with vertical bar indicating the 95% confidence interval. N=number of measurements. p-value derived from a two-sided Kolmogoroff-Smirnoff test. **b**, Force propagation between active and passive tether for *mec-2* mutant neurons. Solid green line indicates a fit to the an exponential decay as described in the methods, shaded area indicating the 95% confidence interval of the fit and the dotted black line indicates the fit derived from control data (Fig. 1d). Size of the dots corresponds to the absolute force transmitted to the passive tether. Inset: density of MEC-2 along the axon in cultured neurons as a proxy for membrane obstacles. **c**, Simulation of the tension propagation comparing cells with decreasing obstacle density; increasing permeability *k*. The simulation corresponds to the periodic case. Hence, the value *k* is used to calculate *k*_+_ and *k*_−_. **d**, Tension propagation for neurons treated with cyclodextrin to sequester cholesterol. Solid orange line indicates a fit to the an exponential decay as described in the methods, shaded area indicating the 95% confidence interval of the fit and the dotted black line indicates the fit derived from control data (Fig. 1d). Size of the dots corresponds to the absolute force transmitted to the passive tether.

MEC-2 stomatin is a cholesterol-binding protein, and the disruption of its cholesterol-binding ability alters the regulation of the ion channel [41, 82]. The addition of cholesterol to biomembranes has been shown to increase the membrane expansion modulus (*E*_*m*_), effectively making the membranes stiffer [83]. This prompted us to investigate how cholesterol depletion affects tension propagation in cultured neurons. To this end, we treated cells with methyl-*β*-cyclodextrin (mbCD) to extract membrane cholesterol and confirmed depletion by monitoring loss of filipin fluorescence, a dye commonly used to visualize membrane cholesterol (Extended Data Fig. 7a, b). As shown above, stiffer membranes propagate tension better (Extended Data Fig. 4d), consequently we would expect that the addition of cyclodextrin to remove cholesterol lowers tension propagation. However, experiments revealed the opposite: treating neurons with 3 mM cyclodextrin to sequester cholesterol significantly increased tension propagation (control vs. treated: 12 µm vs. 17.6 µm), similar to the effects observed with genetic disruption of *mec-2* (Fig. 5d). This surprising increase in tension propagation is inconsistent with the expected effect of softer membranes alone (Fig. **??**g). Instead, we propose that it results from a reduction in obstacle density and clustering within the membrane, as seen in both *mec-2* mutants and cholesterol-depleted neurons. Indeed, a mutant MEC-2 protein with a conversion of proline to serine (P134S) that cannot bind to cholesterol was shown to disrupt proper localization and organization of MEC-2 in the membrane of touch receptor neurons [41]. Collectively, these data suggest that MEC-2 condensates act as an obstacle, interacting with cholesterol in the membrane to limit the propagation of membrane tension.

### 1.2 Conclusions

Using *C. elegans* as a model for mechanotransduction and neuroscience, we dissected the molecular signature of mechanical stress propagation along the neuronal membrane of sensory neurites in neurons responsible for the sense of touch and proprioception. With an optical-tweezer-based membrane tether extrusion assay, we found that, unlike our previous observations of tubes extruded from zebrafish progenitor cells [45, 3] or the soma of cultured TRN [31, 32], the tension did not remain constant during the extrusion protocol, suggesting a limited membrane reservoir that cannot buffer tension gradients in axons [7]. We found that steep gradients of membrane tension propagate fast (*>*120 µm/s) but only to a limited distance (< 20 µm). We also observed considerable variability in the propagation length scale (Fig. 1d) and velocity (Extended Data Fig. 3e), which may reflect the involvement of distinct propagation mechanisms, such as transmission through the membrane, the cytoskeleton, the viscoelastic cytoplasm, and the density and arrangements of obstacles in the membrane. It may also suggest that obstacles are not homogeneously distributed along the neurite, with varying density in proximal and distal regions, as we observed with MEC-2 in our cultures (Extended Data Fig. 7c).

Our experimental results are consistent with predictions from a biophysical model that a higher permeability (lower obstacle density) through membrane-cortex disruption propagates tension farther. Interestingly, we observed a significant dose-dependent effect when altering the actin and microtubule cytoskeleton and spectrin organization. For example, actin depolymerization with varying concentrations of LatA and CytoD had opposing effects on tension propagation. At acute delivery of low concentrations, the propagation distance increased, presumably due to the transient decoupling of membrane cortex binders, while at high and chronic treatments, tension propagation was reduced. Therefore, we speculate that chronic actin depolymerization disrupted caveolae and other membrane domains, essentially increasing the available membrane lipid reservoir as proposed also previously [84, 85, 61]. Indeed, our collective results show that there is no strong correlation between the extent of membrane tension propagation, velocity and resting tension, indicating that each property is regulated by different mechanisms.

We uncovered that periodic obstacle arrangement supports tension propagation over short distances, whereas more homogeneous and randomly organized obstacles better support long-range tension propagation. We propose that mechanical signals must remain confined to be locally processed, much like local computations at synapses. Why might restricted propagation be advantageous? One possibility is that it enhances the cell’s ability to detect precise, localized mechanical cues, enabling parallel processing of simultaneous stimuli and targeted responses. Allowing unrestricted propagation of mechanical information along the axon might blur receptive field boundaries, introducing disruptive background noise. Our data showing that tension propagation is different in touch receptors and proprioceptors suggests differential expression of membrane obstacles such as cortex crosslinkers. Recent single cell gene expression data sets show that DVA expresses 10 times more of the FERM domain protein-1 (FRM-1) than TRNs[86], the major Band 4.1N ortholog of the FERM (Four-point-one, Ezrin, Radixin, Moesin) superfamily in *C. elegans* with roles in coupling various transmembrane receptors, channels, and adhesion molecules to actin /spectrin cytoskeleton or stomatin proteins[87]. In fact, we have recently shown that FRM-1 is directly bound to UNC-70[79], but future work is necessary to refine the role of FRM-1 in this process.

## 2 Acknowledgements

We would like to thank the NMSB laboratory for discussion, Santiago Ortiz-Vásquez (NMSB), Martín Fernández-Campo (NMSB) and Arnau Farré (IMPETUX) for time-shared dual trap characterization, Felix Campelo (ICFO) for suggestions early in the work, Fred Hoerndli (Colorado State) for comments on the draft and the *Caenorhabditis elegans* center (CGC, supported by the National Institutes of Health, P40 OD010440) for reagents.

MK acknowledges financial support from the ERC (MechanoSystems, 715243), Human Frontiers Science Program (RGP021/2023), MCIN/AEI/10.13039/501100011033/FEDER “A way to make Europe” (PID2021-123812OB-I00, CNS2022-135906), “Severo Ochoa” program for Centres of Excellence in R&D (CEX2019-000910-S), from Fundació Privada Cellex, Fundació Mir-Puig, and from Generalitat de Catalunya through the CERCA and Research program. ICFO is the recipient of the Severo Ochoa Award of Excellence of MINECO (Government of Spain). This work was supported in part by 1RF1DA055668-01, 5R01 MH139350 and the Air Force Office of Scientific Research Grant FA955018-1-0051 to P.R.

## 3 Author contribution

Experiments: FCC, NSC. Simulations: MBQ. Analysis: FCC, MBQ, MK. First draft: MK, FCC. Supervision: PR, MK. Funding: PR, MK. All authors edited and revised the manuscript.

## 6 Materials and Methods

### 6.1 *C. elegans* strains and maintenance

Strains were maintained and manipulated under standard conditions [88]. All strains used in this study are enumerated in Supplementary Table S2. Nematode strains were grown at 20°C on nematode growth medium (NGM) plates with OP50 bacteria and synchronized using the standard alkaline hypochlorite treatment method [89].

### 6.2 Primary cell culture and membrane extrusion

#### 6.2.1 TRN primary cell culture

Primary cell culture was performed as described in [31, 10]. In short, gravid hermaphrodites grown onto large, peptone-enriched plates seeded with *E. coli* NA22 bacteria at 20-25°C were collected with milli-Q H2O and bleached following regular procedures [89]. Eggs were washed 3x with Egg Buffer (118 mM NaCl, 48 mM KCl, 2 mM CaCl_2_, 2 mM MgCl_2_, 25 mM HEPES, pH 7.3 and 340 mOsm) and separated through a final sucrose density gradient 30 % generated by 20 min centrifugation at 1300 × g. Once the eggs were collected in a new vial, they were washed 2x with Egg Buffer and treated with chitinase (0.5 U/ml) for 40 min to digest their egg shell. The reaction was stopped with L15 medium. The embryos were then dissociated by passing the solution 10x through a 25G needle and filtered with a 5 µm Durapore filter (Millipore). Finally, cells were pelleted by 3 min centrifugation at 400 xg and resuspended with L15 medium. Cells were seeded at density 1.5 · 10^6^*mL*^−1^ in optical trapping chambers (see below) and 1 ml of L15 medium was added after 3 h. They were cultured at 25°C and fresh medium was changed (1-2 mL) every day. All experiments were performed between two and four days after isolation. CO_2_-independent L15 tissue culture medium supplemented with heat-inactivated bovine calf serum (10% v/v; FBS 11A Capricorn Scientific), penicillin (10 mg mL-1), and streptomycin (100 units; L0022 Biowest) was used throughout the study[31].

The drugs were added 24-48 hours after isolation. Nocodazole (M1404, Sigma) was added at a final concentration of 1 µg/mL. Cytochalasin D (10 mM stock in DMSO, C2618, Sigma) was diluted 1:5000 in L15 medium to 2 µM final concentration. Latrunculin A (10 mM stock, L5163 Sigma) was diluted in L15 medium to a final concentration of 500 nM, 1 µM or 10 µM. Cyclodextrin (Methyl-*β*-cyclodextrin C4555, Sigma) was added to the final concentration of 3 or 10 mM. Cells were incubated with Nocodazole, Cyclodextrin, or latrunculin A (acute) for 30 min, while treatments with cytochalasin D and latrunculin A were incubated overnight at 25 °C, followed by 3 washes of L15 with the corresponding drug. Hypoosmotic shocks were perfomed by diluting 1:1 L15 medium in water.

### 6.3 Cholesterol staining with filipin

Cells were treated with 10 mM Cyclodextrin 48 h post-isolation, as described above. After a 30 min incubation, both the treated and untreated (control) cells were washed three times with PBS. Cells were then fixed with 4% paraformaldehyde (in PBS) for 15 minutes at room temperature, followed by three additional PBS washes. Filipin III (Fermentek, FLP-001) was added at a final concentration of 0.05 mg/mL in PBS, prepared from a 5 mg/mL stock solution in DMSO, as described[90]. After a 2 h incubation in the dark (note that filipin is highly photosensitive), the samples were washed x3 with PBS. Imaging was performed on a TCS SP8 microscope (Leica Microsystems GmbH, Germany), equipped with an HC PL APO CS2 60x/1.4 oil immersion objective and a hybrid detector (HyD). Images were acquired at 16-bit depth. Filipin III was excited at 405 nm (65.3 µW at the sample plane; 40% transmittance) and fluorescence emission was collected in the 410-465 nm range. For the analysis of the filipin fluorescence intensity, cells were segmented using the trainable Weka segmentation plugin in ImageJ/Fiji. This machine-learning-based tool was trained to distinguish cell regions from the background using manually annotated training data. The resulting segmentation masks were applied to identify individual cells as regions of interest (ROIs). Total fluorescence intensity within each ROI was measured, and background signal was subtracted by quantifying fluorescence in cell-free regions. Statistics were derived from two-tailed unpaired Student’s t-test.

### 6.4 Optical micromanipulation and confocal microscopy

A multimodal platform was used that combined spinning-disk confocal microscopy and optical micromanipulation. Briefly, an optical micromanipulation unit (SENSOCELL, IMPETUX) modulated with a pair of acoustooptic defletors (AOD) was coupled to the rear epifluorescence port of an inverted research microscope (Nikon Ti2 Eclipse). The optical traps were created at the focal plane of a 60x/1.2 water-immersion microscope objective (Nikon). A spinning-disk confocal unit (Dragonfly, Andor) was coupled to the microscope left port. A detailed description of the setup can be found in recent publications [91].

Optical trapping chambers containing primary culture cells were built in a #1.5H glass bottom plate (GWST-5040, Willco). Dishes were coated with a 50-µm polydimethylsiloxane (PDMS) layer, from which a 1×1 cm cavity was drawn, in which the cells were seeded as in [10]. Before the experiment, cells were gently rinsed with L15 medium three times and the cavity was filled with 50 µL of a solution containing 1-µm diameter polystyrene beads (red fluorescent, carboxylate modified: F8816, Thermofisher, 1:2000 dilution). Finally, the cavity was cautiously closed with a 25Ø25mm cover slip and mounted on the microscope.

All optical force measurements in the article were carried out through calibration-free detection of light momentum changes using a direct force sensor (SENSOCELL, IMPETUX) collecting the light emerging from the optical traps. The nominal trap stiffness was 3 pN/µm/mW (for a polystyrene 2*R* = 1 µm bead). For a typical experiment with a 360 mW trap (3 W of output laser power, 1064 nm), a trapping stiffness of 1080 pN/µm results into a roll-off frequency of *f*_*c*_ = *k/*(2*πγ*) *∼*17.7 kHz [92], where *γ* = 6*π η*(*T*) *R b* is the friction coefficient (*η*(*T*): medium viscosity; *D*: bead diameter; *b*: Faxén hydrodynamic correction for *z* = 2 µm with respect to the coverglass, see Supplementary Material and Ref. [93]). Room temperature around the microscope table was *T*_room_ = 21 °C, while local temperature around the optically trapped beads was estimated as in ref. [94] as *T* = 24.6 °C.

Simultaneous optical trapping of several microbeads was performed by time-sharing the laser spot at 25 kHz. Details of the time-sharing can be found in ref. [23]. In short, the 1064 nm laser beam with power *P* is split into two alternating traps every 40 µs using a pair of acousto-optic defelectors. During the 40 µs trapping period, each bead feels the power *P*, and thus the power with which the bead is trapped remains constant and does not change with the number of the traps. Consequently, the stiffness of the trap remains constant and is independent of the number of the traps. A thorough analysis of the dual-trap time sharing measurement protocol is introduced in the Supplementary Material.

Because the refreshing rate of the two traps is faster than the bead can diffuse away, or is pulled back into the cell, this allows for quasi-simultaneous measurement of membrane tension at different sites of the neuronal axon through the extrusion of membrane nanotubes (see Supplemetary Material). Because *C. elegans* is a poikilotherm organism and thrives between 15 and 25 degrees Celsius, optical trapping was performed at room temperature (20 degrees Celsius) and the dynamic viscosity of L15 supplemented with 10% fetal bovine serum was 0.930 ± 0.034 mPa s as determined before (ref. [95]). Dynamic control of the optical traps was performed with the manufacturer software (LightACE 1.6.2, IMPETUX) and automated with a software development kit (SDK) in LabView (National Instruments).

#### 6.4.1 Extrusion of membrane nanotubes and spatiotemporal force spectroscopy

Once both microspheres were optically trapped, brief contact (∼1 s) with the axonal membrane was sufficient to extrude two lipid nanotubes (Fig. 1a). Maintaining these tethers in a fixed position required a holding force *F*_*rest*_, which reflects the baseline (resting) membrane tension. While trap 2 was held constant, the active trap 1 was moved at 40 µm s^−1^ for 10 µm. The extrusion of a nanotube translated into an increase in the force signal, which reached a peak, *F*_peak_, of increasing height for a higher pulling velocity. Once the extruding bead stopped, the peak force was followed by viscoelastic relaxation, reaching a static tension value *F*_static_ (Fig. 1b). The tether neck from which the membrane nanotube was extruded could slide along the axon as the optically trapped bead was moved parallel to it (Video 1), reaching a perpendicular orientation with respect to the axon to minimize the nanotube length [8]. This allowed us to separate the tether necks apart (distance *d*, Fig. 1c) to measure membrane tension diffusion at increasing distances along the axon.

The friction of the microbead with respect to water was calculated using the Stokes drag expression corrected for the bead-to-surface interaction ([93], Eq. 3):

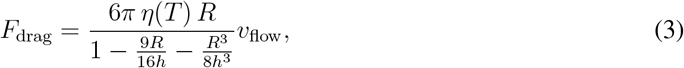

in which *v* is the trap velocity, *h* is the bead-surface separation, *R* is the bead radius and *η*(*T*) is the temperature-dependent viscosity of the medium. The denominator in Eq. 3 accounts for the bead-to-surface hydrodynamic coupling [93]. Heating is negligible because neurons are grown in glass, which has high thermal conductivity[94]. The approach and retraction cycles of a trapped microsphere without attachment to the neuronal membrane did not elicit cellular changes[10].

##### Calculating resting membrane tension

The baseline resting tension of the membrane was determined on the passive and the active tether before each tension propagation experiment (Fig. 1c and Extended Data Fig. 3b). We divided the measured force with the predetermined correction factor *h* to account for the time-sharing deviation (see above). As shown in Extended Data Fig. 2c, when these values come from dual tether extrusion experiments, the correction factor was determined to be 0.63. The resting tension extracted from the active and the passive tethers were not statistically different (Extended Data Fig. 3a) -consequently we pooled these values to obtain the population measure of *σ*_*rest*_. This value depended on the cell type, but also on the pharmacological treatment, and provides information on the mechanical coupling of the membrane to the cortex and the surface tension in the plane[3]. We calculated the effective membrane tension as before[48] using the well-known relation, assuming a bending rigidity of 2.7*·*10^−19^ Nm[43]:

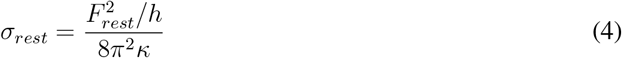

#### 6.4.2 Peak and decay membrane tension dynamics

Membrane tension during tether pull and time relaxation was fitted, respectively, an exponential+linear load function (Eq. 5a) and a stretched exponential decay (Eq. 5b) to both the active and the passive tethers as follows (see Extended Data Fig. 8-insets). A stretched exponential decay captures the multiple time scales arising from molecular heterogeneity [96, 97].

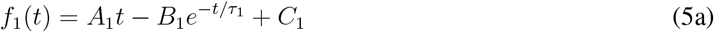

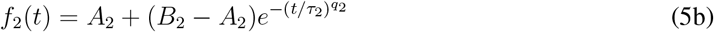

From the fits, we obtained the force peaks for the active and passive tethers as 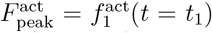 and 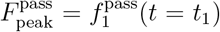, as well as the static tension values after the relaxation, 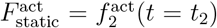 and 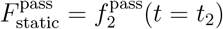. See a graphical explanation in Extended Data Fig. 8.

#### 6.4.3 Measuring force propagation between two tethers along the plasma membrane

The experiments involved an active trap holding a microsphere, which pulled on the corresponding nanotube at a rate of 40 µm/s, while the passive trap remained stationary. The bead was then approached back to the axon after 5 s. The membrane tension gradients originated from the active bead trap were partially diffused into the passive (Fig. 1c). As the tether force at zero velocity reads as 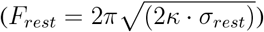 we assume that the change in membrane surface tension at the two tether necks follows Eq. 1[10, 43], from which the relative change in force on the passive trap, with respect to the active, was calculated as in Eq. 6:

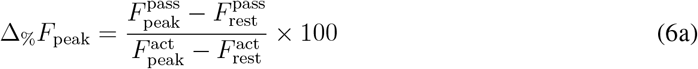

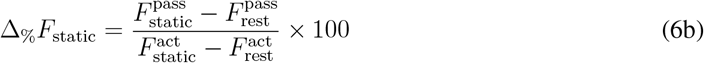

The force transmission (Fig. 1d) reached 40-50% for the shortest double tether separation, *d →* 0. To achieve this, the two beads faced each other and pulled the lipid nanotubes towards opposite directions. For increasing *d*, the tension rapidly dropped from the *active* to the *passive* tether, suggesting effective buffering of the membrane tension along the axon. When 20 µm apart, the tension in the passive nanotube almost vanished (Δ*F*_%_ < 20%).

To determine the best-fitting model while accounting for both fit quality and model complexity, we employed the AIC for model comparison. The AIC is defined *AIC* = 2*k −*2 ln(*L*), where *k* is the number of estimated parameters and *L* is the maximum likelihood of the model. Lower AIC values indicate models that achieve a better trade-off between explanatory power and parsimony. All model fitting and AIC calculations were performed in R using the lm() and AIC() functions. We formally compared three functions, *f* (*x*) = *A* exp(−*x/δ*), *g*(*x*) = *A* exp(−*x/δ*) + *C* and *h*(*x*) = *A* exp(−*x*^2^*/δ*). To quantify the spatial decay of force propagation, we plotted the ratio of the peak force in the passive tether to the peak force in the active tether as a function of the distance between the two tethers (Fig. 1d) and fitted a mono-exponential decay model exp(*x*) = *A* exp(−*x/δ*) using weighted least squares.

Model fitting was performed using the Levenberg–Marquardt algorithm via the nlsLM() function from the minpack.lm R library. To improve robustness against outliers and heteroscedasticity, we implemented a two-step fitting procedure: an initial fit was used to calculate residuals, which were then transformed into observation-level weights using a Huber-type loss function (tuning constant k=1.345; ref. [98]). These weights were then used in a second fit, that was more robust against influential data points. To identify and visualize influential observations, we computed Cook’s distance based on a linearized approximation of the nonlinear model. This was done by fitting a linear model to the same response and predictor variables and calculating the standard leverage-based influence measure using the formula: 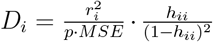 where *r*_*i*_ is the residual for observation *i, h*_*ii*_ is the leverage, *p* the number of parameters, and MSE the mean squared error of the linear fit. Points with *D*_*i*_ *>* 4*n* were flagged as potentially influential and highlighted in the final visualization.

The variance–covariance matrix of parameter estimates was extracted from the fitted model, and 95% confidence intervals on the fitted curves were computed by propagating uncertainty via the Jacobian matrix of the model at prediction points. Notably, nlsLM yielded narrower 95% confidence intervals compared to nls (which uses Gauss-Newton algorithm), likely due to improved numerical stability and better conditioning of the Jacobian used in estimating parameter uncertainty. The final plot displays the robust fits and confidence ribbons for both control and test groups across the measured spatial domain.

To account for measurement uncertainties in offset compensation during the calibration procedure [91], the initial fit was weighted using the absolute peak force of the passive tether.

#### 6.4.4 Measuring the tension propagation speed from the active to the passive tether

The time delay between the two peaks was measured by identifying a maximum in the correlation between the active and passive tether forces, which can be expressed as:

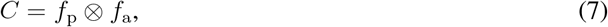

where *⊗* represents the correlation operation and was carried out using the *xcorr* function in Matlab (version R2019b). We estimated the mean propagation velocity in each condition by fitting a linear regression of distance (*d*) on delay (Δ*t*), *d* = *ν*Δ*t* + *b*, so that the slope *ν* is directly the velocity. Because a power-law model with exponent 0.5 yielded a lower AIC (593) than the linear model (604), we applied a square transformation to linearize the relationship. To test for statistically significant differences in velocity between treatments, we then modeled the squared distance as a function of delay, *d*^2^ = *β*_0_ + *β*_1_ · Δ*t*, and compared the *β*_1_ coefficients (with their confidence intervals) across treatment groups as described below.

### 6.5 Tension propagation model

Following Shi *et al*. [15], we augment the Stokes equation with a drag term 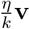**v**, which gives

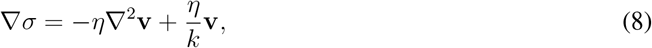

where *σ* is the tension, **v** is the velocity field of the lipid flow, *η* is the two-dimensional membrane viscosity, *k* is the Darcy permeability array of the obstacles. We also assume conservation of mass, given by

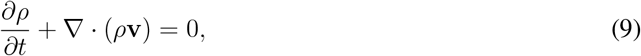

where *ρ* is the two-dimensional density of lipids. As in [15], we assume a small perturbationin the lipid density *ρ*_0_ + *ϵρ*, then *∇·* ((*ρ*_0_ + *ϵρ*) **v**) = *∇* (*ρ*_0_ + *ϵρ*) *·* **v** + (*ρ*_0_ + *ϵρ*) *∇·* **v** = *ρ*_0_ *∇·* **v** + *∇·*(*ϵρ***v**). Thus, a change in the density *δρ* is given by −*ρ*_0_*∇·***v**. Following [15], we assume a linear stress-strain relationship for the membrane tension *δσ* = −*E*_*m*_*δρ/ρ*_0_, where *E*_*m*_ represents the effective area expansion modulus of the cell membrane. Substituting the change in density, we get *δσ* = *E*_*m*_ *∇·* **v**.

Now, taking the divergence of Eq. 8

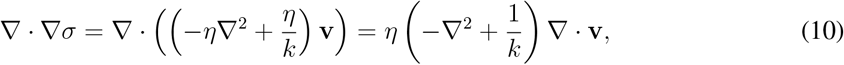

and the above stress-strain relationship gives

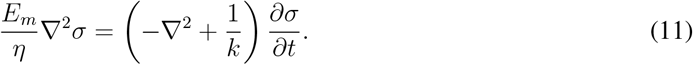

Note that, differently from Shi *et al*. [15], we do not assume that the spacing between transmembrane obstacles is small compared to externally imposed variations in the flow field. This allows us to investigate different spatial configurations of the obstacles.

We solve Eq. (11) in COMSOL Multiphysics 6.1 using the parameters in Supplementary Table 3. The pulling events were simulated by combining the definition of the tension in the tether (*σ*_*t*_),

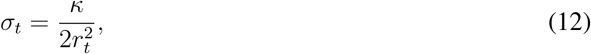

with the area of the tether

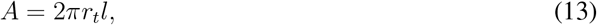

which gives

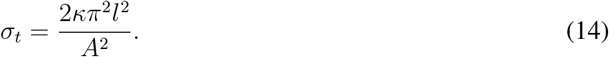

Here, *κ* is the bending stiffness of the membrane, *r*_*t*_ is the tether radius, and *l* is its length. Thus, the change of the tension due to the pulling of the tether is given by

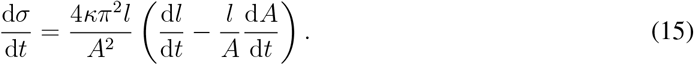

We take 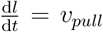 and 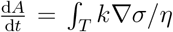 where *T* is the perimeter of the tether boundary and *v*_*pull*_ the pulling velocity. The tension propagation curves in Fig. **??** were obtained by measuring the tension at the tip of the active and passive tether. Note that we assume an initial length of the tether *l*_0_ and did not change the geometry of the model during the simulation. Instead, we include the tension flow at the tip of the tether corresponding to Eq. 15.

### 6.6 Statistics and reproducibility

All statistical calculations were performed in R (version 4.2.2, 2022-10-31), Python (version 3) or Matlab. The Cliff’s *delta* was chosen as a non-parametric effect size measure to quantifies the degree of overlap between two distributions when the data was not normally distributed or when comparing medians instead of means (Table 1). Setting *X* and *Y* as the two groups, we use 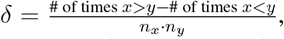,where *n*_*x*_ and *n*_*y*_ are the sample sizes of groups *X* and *Y*, respectively. The result is between -1 and 1 and 1 means all values in group *X >* group *Y*, −1 means all values in group *X <* group *Y*, and 0 means complete overlap (no effect).

For standard hypothesis testing, we exclusively performed two-sided tests, unless otherwise indicated in the figure legend. The level of significance for all comparison was chosen at *α*=0.05, unless otherwise indicated in the figure or figure legend. No statistical methods were used to pre-determine sample sizes but our sample sizes are similar to those reported in previous publications. Sample sizes are indicated in the figures. Data collection and analysis were not performed blind to the conditions of the experiments, except when indicated otherwise. Unless otherwise stated, all data have been assumed to not follow a normal distribution. Accordingly, distributions of the baseline resting tensions (*σ*_*rest*_, e.g. Fig. 1d) were tested using a Kolmogoroff-Smirnoff test for pairwise comparison using wild-type as a control group. The 95% confidence interval was derived by bootstrapping, in which we randomly sample N data points with replacements from the measured distribution, where N is the sample size of the experimental data set. This procedure was repeated 1000 times and the median was calculated each time. Finally, we sort the bootstrapped distribution of medians and select the 2.5th and 97.5th percentiles for the 95% CI.

Only mono or bi-polar, healthy looking neurons were used for the experiment, and cells not clearly showing a differentiated axon/neurite, or with extensive neurite branches were not used for measurements. As some treatments have previously been shown to result in aberrant neurite morphologies and branches, this may introduce a bias towards ‘wt’-like neurons, but avoids confounding effects due to geometry (large diameter varicosities, branches) and cell-substrate contacts. Tether extrusions with more than one attachment between the axon and the bead were excluded from the analysis.

To assess differences in force transmission between experimental and control conditions, we analyzed the ratio of transmitted to active force as a function of distance. The ratio was first log-transformed to normalize the distribution and stabilize variance. A linear regression was then performed with distance as the independent variable and the log-transformed force ratio as the dependent variable.

To account for variability in measurement confidence, the regression was weighted by the absolute value of the transmitted force. This approach reflects the assumption that lower transmitted forces are associated with increased uncertainty, and therefore should contribute less to the regression model. Comparisons between experimental and control conditions were made by evaluating the difference in regression slopes, testing whether the spatial distribution of force transmission varied significantly across groups. The resulting p-value quantified the statistical significance of this difference. All analyses were carried out in R, using weighted linear models between *x* (distance) and *y* (the force ratio):

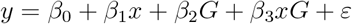

where *G* is a binary indicator variable for group (0 = control, 1 = test), and is the residual error, *β*_0_ is the intercept for the control group, *β*_1_ is the slope for the control group,*β*_2_ represents the difference in intercept between groups, and *β*_3_ represents the difference in slope between groups. A statistically significant interaction term (*β*_3_, p < 0.05) indicates that the slopes of the regression lines differ significantly between groups. Model fitting was performed in R using the lm() function and statistical tests were based on a t-test according 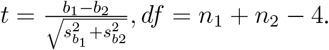.

#### 6.6.1 Simulations

Similar to the experiments, we calculated the extent of tension propagation after plotting the ratio of the peak tension in the passive tether to the peak tension in the active tether as a function of the distance between the two tethers (Figs. **??**c, Extended Data Fig. 4) and fitted an exponential decay function exp(*x*) = *A* exp(−*δx*) using weighted least squares. We used the Matlab function fit.m, and to evaluate the fit, we obtained the root mean squared error. We noticed a systematic underestimation of the exponential fit function for large distances between tethers in the simulations. While force transmission dissipates for long distances in experiments, we found a residual tension for our given set of parameters in the model simulations, which cannot be captured by the exponential fit. The parameter choices for the model were based on: 1) having a physiologically feasible set of parameters and 2) replicating the levels of tension propagation at small distances. Thus, instead of fitting the parameters to the experimental data, we decided to keep the physiologically feasible parameters in the model and attributed the residual tension at long distances to effects that are not accounted for in the model, such as sources and sinks of lipids via endo-and exocytosis. We acknowledge that future iterations of the model should address this discrepancy. To better model the extent of tension propagation, we incoporated an offset *C* to the exponential function, exp(*x*) = *A* exp(−*δx*) + *C*. This addition resulted in better fittings, for *k* = 0.03 µm^2^ AIC decreased from 42.99 to 8.24 in the random case and from 48.95 to 11.06 in the periodic case (the fittings with the offset are in Figure **??**). Hence, the exponential decay allowed us to better compare different conditions in the simulations. To test if the difference between conditions of the extent of tension propagation *δ* was significant, we obtained the confidence interval with the confint.m function and obtained the standard error (SE) with Student’s t inverse cumulative distribution function, tinv.m. Then, we calculated the t-score for a two-tailed t-test by taking 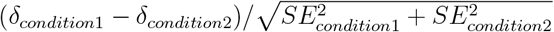.

## 7 Data availability

All numerical data are supplied as a source data sheet in the Supplementary Material of this manuscript. All raw data have been deposited on zenodo.org under doi: 10.5281/zenodo.15465212

## 8 Code availability

Code for tension propagation measurements is supplied through the Gitlab account here. Code to perform the timesharing optical trapping simulations is supplied through this link.

## References

1. Ghisleni, A., Gauthier, N. C., Mechanotransduction through membrane tension: It’s all about propagation? Current Opinion in Cell Biology 86 (2024).

2. Roux, A. L. L., Quiroga, X., Walani, N., Arroyo, M., Roca-Cusachs, P., The plasma membrane as a mechanochemical transducer. Philosophical Transactions of the Royal Society B: Biological Sciences 374 (2019).

3. Diz-Muñoz, A., et al., Control of directed cell migration in vivo by membrane-to-cortex attachment. PLoS Biology 8, e1000544 (2010).

4. Sukharev, S., Sachs, F., Molecular force transduction by ion channels: diversity and unifying principles. Journal of Cell Science 125, 3075–3083 (2012).

5. Appadurai, D., et al., Plasma membrane tension regulates eisosome structure and function. Molecular Biology of the Cell 31, 287–303 (2020).

6. Dai, J., Sheetz, M. P., Cell Membrane Mechanics, vol. 55, 157–177. Academic Press (1998).

7. Raucher, D., Sheetz, M. P., Characteristics of a membrane reservoir buffering membrane tension. Biophys J 77, 1992–2002 (1999).

8. Datar, A., Bornschlögl, T., Bassereau, P., Prost, J., Pullarkat, P. A., Dynamics of membrane tethers reveal novel aspects of cytoskeleton-membrane interactions in axons. Biophysical Journal 108, 489–497 (2015).

9. Brownell, W. E., Qian, F., Anvari, B., Cell membrane tethers generate mechanical force in response to electrical stimulation. Biophysical Journal 99, 845–852 (2010).

10. Das, R., et al., An asymmetric mechanical code ciphers curvature-dependent proprioceptor activity. Science Advances 7, 1–20 (2021).

11. Pradhan, S., Williams, M. A., Hale, T. K., Changes in the properties of membrane tethers in response to hp1α depletion in mcf7 cells. Biochemical and Biophysical Research Communications 587, 126–130 (2022).

12. Dai, J., Sheetz, M. P., Wan, X., Morris, C. E., Membrane tension in swelling and shrinking molluscan neurons. J Neurosci 18, 6681–6692 (1998).

13. Rangamani, P., The many faces of membrane tension: Challenges across systems and scales. Biochimica et Biophysica Acta - Biomembranes 1864 (2022).

14. Mashanov, G. I., et al., Heterogeneity of cell membrane structure studied by single molecule tracking. Faraday Discussions 232, 358–374 (2021).

15. Shi, Z., Graber, Z. T., Baumgart, T., Stone, H. A., Cohen, A. E., Cell membranes resist flow. Cell 175, 1769–1779.e13 (2018).

16. Rangamani, P., Mandadap, K. K., Oster, G., Protein-induced membrane curvature alters local membrane tension. Biophysical Journal 107, 751–762 (2014).

17. Belly, H. D., et al., Cell protrusions and contractions generate long-range membrane tension propagation. Cell 186, 3049–3061.e15 (2023).

18. Shi, Z., Innes-Gold, S., Cohen, A. E., Membrane tension propagation couples axon growth and collateral branching. Science Advances 8 (2022).

19. Perez, C. G., et al., Rapid propagation of membrane tension at a presynaptic terminal. Science Advances 4411, 2021.05.26.445801 (2022).

20. Jauffred, L., Callisen, T. H., Oddershede, L. B., Visco-elastic membrane tethers extracted from escherichia coli by optical tweezers. Biophys J 93, 4068–4075 (2007).

21. Guilford, W. H., Tournas, J. A., Dascalu, D., Watson, D. S., Creating multiple time-shared laser traps with simultaneous displacement detection using digital signal processing hardware. Analytical Biochemistry 326, 153–166 (2004).

22. Capitanio, M., Cicchi, R., Pavone, F. S., Continuous and time-shared multiple optical tweezers for the study of single motor proteins. Optics and Lasers in Engineering 45, 450–457 (2007).

23. Català-Castro, F., et al., Measuring age-dependent viscoelasticity of organelles, cells and organisms with time-shared optical tweezer microrheology. Nature Nanotechnology (2025).

24. Sanzeni, A., et al., Somatosensory neurons integrate the geometry of skin deformation and mechanotransduction channels to shape touch sensing. eLife 8, 1–44 (2019).

25. Krieg, M., Dunn, A. R., Goodman, M. B., Mechanical systems biology of c. elegans touch sensation. BioEssays 37, 335–344 (2015).

26. Krieg, M., et al., Genetic defects in β-spectrin and tau sensitize c. elegans axons to movementinduced damage via torque-tension coupling. eLife 6, e20172 (2017).

27. Das, A., et al., C. elegans touch receptor neurons direct mechanosensory complex organization via repurposing conserved basal lamina proteins. Current Biology 1–19 (2024).

28. Cueva, J. G., Mulholland, A., Goodman, M. B., Nanoscale organization of the mec-4 deg/enac sensory mechanotransduction channel in caenorhabditis elegans touch receptor neurons. Journal of Neuroscience 27, 14089–14098 (2007).

29. Sanfeliu-Cerdán, N., et al., A mec-2/stomatin condensate liquid-to-solid phase transition controls neuronal mechanotransduction during touch sensing. Nature Cell Biology 25, 1590–1599 (2023).

30. O’Hagan, R., Chalfie, M., Goodman, M. B., The mec-4 deg/enac channel of caenorhabditis elegans touch receptor neurons transduces mechanical signals. Nature Neuroscience 8, 43–50 (2005).

31. Krieg, M., Dunn, A. R., Goodman, M. B., Mechanical control of the sense of touch by β-spectrin. Nature Cell Biology 16, 224–233 (2014).

32. Vásquez, V., Krieg, M., Lockhead, D., Goodman, M. B., Phospholipids that contain polyunsaturated fatty acids enhance neuronal cell mechanics and touch sensation. Cell Reports 6, 70–80 (2014).

33. Bennett, V., Healy, J., Membrane domains based on ankyrin and spectrin associated with cell-cell interactions. Cold Spring Harbor perspectives in biology 1 (2009).

34. Xu, K., Zhong, G., Zhuang, X., Actin, spectrin, and associated proteins form a periodic cytoskeletal structure in axons. Science (New York, NY) 339, 452–456 (2013).

35. Deng, H., et al., Spectrin couples cell shape, cortical tension, and hippo signaling in retinal epithelial morphogenesis. Journal of Cell Biology 219 (2020).

36. Fletcher, G. C., et al., Mechanical strain regulates the hippo pathway in *drosophila*. Development dev.159467 (2018).

37. Dubey, S., et al., The axonal actin-spectrin lattice acts as a tension buffering shock absorber. ELife 9, 1–22 (2020).

38. Mylvaganam, S., et al., The spectrin cytoskeleton integrates endothelial mechanoresponses. Nature Cell Biology 24, 1226–1238 (2022).

39. Lorenzo, D. N., et al., βii-spectrin promotes mouse brain connectivity through stabilizing axonal plasma membranes and enabling axonal organelle transport. Proceedings of the National Academy of Sciences 116, 15686–15695 (2019).

40. Qi, Y., et al., Membrane stiffening by stoml3 facilitates mechanosensation in sensory neurons. Nature Communications 6, 8512 (2015).

41. Huber, T. B., et al., Podocin and mec-2 bind cholesterol to regulate the activity of associated ion channels. Proceedings of the National Academy of Sciences of the United States of America 103, 17079–17086 (2006).

42. Bussell, S. J., Koch, D. L., Hammer, D. A., Effect of hydrodynamic interactions on the diffusion of integral membrane proteins: diffusion in plasma membranes. Biophysical Journal 68, 1836–1849 (1995).

43. Hochmuth, R. M., Shao, J. Y., Dai, J., Sheetz, M. P., Deformation and flow of membrane into tethers extracted from neuronal growth cones. Biophys J 70, 358–369 (1996).

44. Li, Z., et al., Membrane tether formation from outer hair cells with optical tweezers. Biophys J 82, 1386–1395 (2002).

45. Krieg, M., Helenius, J., Heisenberg, C. P., Muller, D. J., A bond for a lifetime: Employing membrane nanotubes from living cells to determine receptor-ligand kinetics. Angewandte Chemie International Edition 47, 9775–9777 (2008).

46. Borghi, N., Brochard-Wyart, F., Tether extrusion from red blood cells: integral proteins unbinding from cytoskeleton. Biophys J 93, 1369–1379 (2007).

47. Bar-Ziv, R., Moses, E., Instability and “pearling” states produced in tubular membranes by competition of curvature and tension. Physical Review Letters 73, 1392–1395 (1994).

48. Sheetz, M. P., Cell control by membrane-cytoskeleton adhesion. Nat Rev Mol Cell Biol 2, 392–396 (2001).

49. Li, W., Feng, Z., Sternberg, P. W., A c. elegans stretch receptor neuron revealed by a mechanosensitive trp channel homologue. Nature 440, 684–687 (2006).

50. Argudo, D., Capponi, S., Bethel, N. P., Grabe, M., A multiscale model of mechanotransduction by the ankyrin chains of the nompc channel. Journal of General Physiology 151, 316–327 (2019).

51. Wang, Y., et al., The push-to-open mechanism of the tethered mechanosensitive ion channel nompc. eLife 10, 1–20 (2021).

52. Brohawn, S. G., Su, Z., MacKinnon, R., Mechanosensitivity is mediated directly by the lipid membrane in traak and trek1 k+ channels. PNAS 111, 3614–3619 (2014).

53. Krieg, M., Pidde, A., Das, R., Mechanosensitive body–brain interactions in caenorhabditis elegans. Current Opinion in Neurobiology 75, 102574 (2022).

54. Katta, S., Krieg, M., Goodman, M. B., Feeling force: Physical and physiological principles enabling sensory mechanotransduction. Annual Review of Cell and Developmental Biology 31, 347– 371 (2015).

55. Eastwood, A. L., et al., Tissue mechanics govern the rapidly adapting and symmetrical response to touch. Proceedings of the National Academy of Sciences 113, E2471–E2471 (2015).

56. Nekimken, A. L., et al., Pneumatic stimulation of c. elegans mechanoreceptor neurons in a microfluidic trap. Lab Chip 17, 1116–1127 (2017).

57. D’Este, E., Kamin, D., Göttfert, F., El-Hady, A., Hell, S. W., Sted nanoscopy reveals the ubiquity of subcortical cytoskeleton periodicity in living neurons. Cell Reports 10, 1246–1251 (2015).

58. Coste, B., et al., Piezo1 and piezo2 are essential components of distinct mechanically activated cation channels. Science 330, 55 (2010).

59. Kahn-Kirby, A. H., et al., Specific polyunsaturated fatty acids drive trpv-dependent sensory signaling in vivo. Cell 119, 889–900 (2004).

60. Else, P. L., The highly unnatural fatty acid profile of cells in culture. Progress in Lipid Research 77 (2020).

61. Dharan, R., et al., Intracellular pressure controls the propagation of tension in crumpled cell membranes. Nature Communications 16, 91 (2025).

62. Rentsch, J., et al., Sub-membrane actin rings compartmentalize the plasma membrane. Journal of Cell Biology 223 (2024).

63. Spector, I., Shochet, N., Kashman, Y., Groweiss, A., Latrunculins: Novel marine toxins that disrupt microfilament organization in cultured cells. Science 219, 493–495 (1983).

64. Schliwa, M., Action of cytochalasin d on cytoskeletal networks. Journal Cell Biology 92, 79–1 (1982).

65. Qu, Y., Hahn, I., Webb, S. E., Pearce, S. P., Prokop, A., Periodic actin structures in neuronal axons are required to maintain microtubules. Molecular Biology of the Cell 28, 296–308 (2017).

66. Stevenson, B. R., Begg, D. A., Concentration-dependent effects of cytochalasin d on tight junctions and actin filaments in mdck epithelial cells. Journal of cell science 367–375 (1994).

67. Liang, X., Madrid, J., Howard, J., The microtubule-based cytoskeleton is a component of a mechanical signaling pathway in fly campaniform receptors. Biophysical journal 107, 2767–74 (2014).

68. Burke, S. D., Jordan, J., Harrison, D. G., Karumanchi, S. A., Solving baroreceptor mystery: Role of piezo ion channels. Journal of the American Society of Nephrology 30, 911–913 (2019).

69. Morley, S. J.,, Y. Q., A, H. P., Acetylated tubulin is essential for touch sensation in mice. ELife (2016).

70. Chalfie, M., Thomson, J. N., Structural and functional diversity in the neuronal microtubules of caenorhabditis elegans. The Journal of Cell Biology 93, 15–23 (1982).

71. Lockhead, D., et al., The tubulin repertoire of c. elegans sensory neurons and its context-dependent role in process outgrowth. Molecular biology of the cell 27, 3717–3728 (2016).

72. He, L., et al., Cortical anchoring of the microtubule cytoskeleton is essential for neuron polarity. eLife 9, 1–32 (2020).

73. Chalfie, M., Au, M., Genetic control of differentiation of the caenorhabditis elegans touch receptor neurons. Science (New York, N.Y.) 243, 1027–33 (1989).

74. Liu, B. P., Chrzanowska-Wodnicka, M., Burridge, K., Microtubule depolymerization induces stress fibers, focal adhesions, and dna synthesis via the gtp-binding protein rho. Cell Adhesion and Communication 5, 249–255 (1998).

75. Edidin, M., Kuo, S. C., Sheetz, M. P., Lateral movements of membrane glycoproteins restricted by dynamic cytoplasmic barriers. Science 254, 1379–1382 (1991).

76. Bennett, J. S., et al., Spatially-resolved rotational microrheology with an optically-trapped sphere. Scientific Reports 3, 1–5 (2013).

77. Garrido, J. J., et al., A targeting motif involved in sodium channel clustering at the axonal initial segment. Science 300, 2091–2094 (2003).

78. Baines, A. J., Evolution of spectrin function in cytoskeletal and membrane networks. Biochemical Society Transactions 37, 796 (2009).

79. Malaiwong, N., et al., Mechanical load conditions the spectrin network to ‘runon’ proteolysis and promotes early onset neurodegeneration. BioRxiv (2024).

80. Ghisleni, A., Bonilla-Quintana, M., Crestani, M., Rangamani, P., Gauthier, N. C., Mechanically induced conformational transition of spectrin in the mammalian cell cortex. Nature Communications 122, 263a (2024).

81. Wernert, F., et al., The actin-spectrin submembrane scaffold restricts endocytosis along proximal axons. Science (New York, N.Y.) 385, eado2032 (2024).

82. Brown, A. L., Liao, Z., Goodman, M. B., Mec-2 and mec-6 in the caenorhabditis elegans sensory mechanotransduction complex: Auxiliary subunits that enable channel activity. Journal of General Physiology 131, 605–616 (2008).

83. Needham, D., Nunn, R. S., Elastic deformation and failure of lipid bilayer membranes containing cholesterol. Biophysical Journal 58, 997–1009 (1990).

84. Hochmuth, R. M., Micropipette aspiration of living cells. Journal of Biomechanics 33, 15–22 (2000).

85. Echarri, A., Pozo, M. A. D., Caveolae - mechanosensitive membrane invaginations linked to actin filaments. Journal of Cell Science 128, 2747–2758 (2015).

86. Taylor, S. R., et al., Molecular topography of an entire nervous system. Cell 184, 1–19 (2021).

87. Yang, Q., Liu, J., Wang, Z., 4.1n-mediated interactions and functions in nerve system and cancer. Frontiers in Molecular Biosciences 8 (2021).

## Methods only References

88. Stiernagle, T., Maintenance of c. elegans. WormBook : the online review of C. elegans biology 1–11 (2006).

89. de-la Riva, M.P., Fontrodona, L., Villanueva, A., Cerón, J., Basic caenorhabditis elegans methods: Synchronization and observation. Journal of Visualized Experiments e4019 (2012).

90. Biswas, A., Kashyap, P., Datta, S., Sengupta, T., Sinha, B., Cholesterol depletion by mβcd enhances cell membrane tension and its variations-reducing integrity. Biophysical Journal 116, 1456–1468 (2019).

91. Català-Castro, F., Venturini, V., Ortiz-Vásquez, S., Ruprecht, V., Krieg, M., Direct force measurements of subcellular mechanics in confinement using optical tweezers. Journal of Visualized Experiments 2021, 1–35 (2021).

92. Berg-Sørensen, K., Flyvbjerg, H., Power spectrum analysis for optical tweezers. Review of Scientific Instruments 75, 594–612 (2004).

93. Schäffer, E., Nørrelykke, S. F., Howard, J., Surface forces and drag coefficients of microspheres near a plane surface measured with optical tweezers. Langmuir 23, 3654–3665 (2007).

94. Català, F., Marsà, F., Montes-Usategui, M., Farré, A., Martín-Badosa, E., Influence of experimental parameters on the laser heating of an optical trap. Scientific Reports 7, 1–9 (2017).

95. Poon, C., Measuring the density and viscosity of culture media for optimized computational fluid dynamics analysis of in vitro devices. Journal of the Mechanical Behavior of Biomedical Materials 126 (2022).

96. Lukichev, A., Physical meaning of the stretched exponential kohlrausch function. Physics Letters, Section A: General, Atomic and Solid State Physics 383, 2983–2987 (2019).

97. Bonfanti, A., Kaplan, J. L., Charras, G., Kabla, A., Fractional viscoelastic models for power-law materials. Soft Matter 16, 6002–6020 (2020).

98. Huber, P., Robust estimation of a location parameter. Annals Mathematical Statistics 35, 73–101 (1964).

